# High-Throughput Quantification of Population Dynamics using Luminescence

**DOI:** 10.1101/2025.08.26.672247

**Authors:** Malte Muetter, Daniel Angst, Roland Regoes, Sebastian Bonhoeffer

**Affiliations:** Department of Environmental Systems Science, ETH Zürich, Universitätstrasse 16, 8092 Zürich, Switzerland

**Keywords:** bioluminescence, colony forming units, pharmacodynamics, antibiotics, time-kill curves

## Abstract

The dynamics of bacterial population decline at antibiotic concentrations above the minimum inhibitory concentration (MIC) remain poorly characterized. This is because measuring colony-forming units (CFU), the standard assay to quantify inhibition, is slow, labour-intensive, costly, and can be unreliable at high drug concentrations. Luminescence assays are widely used to quantify population dynamics at subinhibitory concentrations, yet their limitations and reliability at super-MIC concentrations remain underexplored. To fill this gap, we compared luminescence- and CFU-based rates across 20 antimicrobials. In our experiments luminescence- and CFU-based rates did not differ significantly for half of them. For the other half, CFU-based estimates of rates of decline were consistently higher. The estimates differed for two main reasons: First, because light intensity tracks biomass more closely than population size, luminescence declined more slowly than the population when bacteria filamented. Second, CFU-based estimates indicated a steeper decline when antimicrobial treatment reduced the number of colonies formed per plated bacterium. This effect can result from changes in clustering behaviour, physiological changes that impair culturability, or antimicrobial carry-over. Thus, the suitability of luminescence to quantify bacterial decline depends on the physiological effects of the antimicrobial used (e.g. filamentation) and whether the quantity of interest is cell number or biomass. Within these limitations, luminescence can serve as an efficient, high-throughput alternative for quantifying bacterial dynamics at super-MIC concentrations.

## Introduction

Accurate characterization of changes in population size under treatment is essential for understanding the evolution of antibiotic resistance. Commonly, the effect of treatment on bacterial populations is quantified by pharmacodynamic (PD) curves. PD curves quantify the relationship between drug concentration and the rate of population change (net growth) [1]. These range from no antibiotic, through sub-MIC concentrations that only reduce population growth, to super-MIC concentrations that kill bacteria and lead to a population decline.

The growth parameter most often used in PD curves is the exponential rate of change in living bacteria, *ψ*_*B*_, reflecting both division and death. However, other population properties, such as changes in the number of culturable bacteria or total biomass, may also be relevant, depending on the specific biological question.

In practice, PD curves are fitted to the rate of change of a measured proxy signal such as optical density (OD), colony-forming units (CFU) or bioluminescent light intensity.

OD is a cost-effective approach for real-time, high-throughput monitoring of culture turbidity without sacrificing the population. OD is positively correlated (within a certain range) to cell density. However, since OD cannot distinguish between living and dead cells, this estimate of cell density is only reliable for increasing or stable population sizes, making it unsuitable for quantifying negative rates (kill rates). Similarly, fluorescence is unsuitable for measuring population decline, as cell death does not inactivate the fluorescent proteins.

CFU assays estimate bacterial density by counting the colonies that grow on permissive agar media from plated samples. They remain the gold standard for measuring population size under both sub-MIC and super-MIC conditions and are widely used to quantify pharmacodynamic curves (e.g. [1, 2]).

Luminescence assays measure the light emitted by bioluminescent bacterial cultures and can be used as a proxy to estimate changes in population size. Two main approaches for biological assays are: eukaryotic *luc* systems and the prokaryotic *lux* systems. The *luc* system, derived from eukaryotes such as fireflies, uses an ATP-dependent luciferase that oxidises luciferin to emit light [3]. It was adapted for bacterial reporters by chromosomal integration in *Mycobacterium tuberculosis* [4] and later tested for quantifying antibiotic killing in *Streptococcus gordonii* [5]. However, *luc*-based assays are limited by sensitivity to intracellular ATP, the need for addition of a costly substrate, and luciferin degradation, making continuous measurement in the same culture impractical.

By contrast, the *lux* operon of prokaryotes such as *Photorhabdus luminescens* encodes all components required to sustain the bioluminescence reaction [6, 7]. No external substrate is needed, so light production can be recorded continuously in the same culture, making the *lux* system better suited for high-throughput applications than the luc system. Accordingly, it has been widely used to record growth curves and quantify sub-MIC treatment effects [8–13].

While high-throughput OD and luminescence measurements at sub-MIC concentrations provide valuable insights into drug effects on growth rates, the super-MIC range is clinically more relevant. A comprehensive investigation of super-MIC population dynamics (for example, pharmacodynamics of drug combinations or resistance mutations) using CFU remains impractical, as it is labor-intensive and inherently low-throughput.

Whether lux luminescence can be extended to super-MIC ranges remains unclear, as direct comparisons between CFU- and luminescence-based measurements are scarce and so far limited to only a few drugs [14–16]. Here we evaluate the potential and limitations of bioluminescent bacteria as a high-throughput model system to quantify population-level net growth rates at the clinically most relevant concentrations (super-MIC). For this, we compare changes in light intensity with changes in CFU counts across 20 antimicrobials spanning 11 classes, including penicillins, cephalosporins, carbapenems, polymyxins, quinolones, rifamycins, tetracyclines, amphenicols, folate antagonists, fosfomycin, and antimicrobial peptides. Luminescence- and CFU-based rates aligned for some antimicrobials (e.g., colistin, amoxicillin) but diverged for others (e.g., ciprofloxacin). In this work, we identify the conditions under which they align with the rate of change of population size, and discuss the implications for studying antimicrobial effectiveness across sub-MIC and super-MIC ranges.

## Results

To investigate the validity of luminescence assays as a high-throughput measure for bacterial population size and its change across the entire range of antimicrobial concentrations, we compared this measure to the number of colony-forming units (CFU). Specifically, we tested whether the rates of change in light intensity, *ψ*_*I*_, and CFU, *ψ*_CFU_, agree for various drugs, and if not, we explored the reasons for any discrepancies.

We used a modified luminescence operon *luxCDABE* from *P. luminescens*. To minimize plasmid copy-number effects on the light emitted by a single cell (cell-specific luminosity), we excised the operon from the pCS–*λ* plasmid ([8]) and inserted it into the *Escherichia coli* chromosome.

### Light intensity is proportional to bacterial density

We first evaluated how the observed bioluminescent light intensity, *I*, which represents a fraction *κ* of the total light emitted by the culture, correlates with bacterial density. To this end, we prepared three replicate overnight cultures of bioluminescent *E. coli*, serially diluted them tenfold, and measured the light intensity for each dilution. We found that light intensity increased linearly with bacterial density for signals above approximately 20 rlu (Figure S1; *R*^2^ = 0.987, *n* = 20, *p <* 10^*−*14^), with a proportionality constant 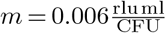. From this observation, we conclude that *κ* remains independent of bacterial density, indicating that, up to one-tenth of the stationary-phase density, high cell densities do not attenuate emitted light. Thus, provided we maintain the same luminescence plate-reader setup, we can assume *κ* to be constant for all subsequent analyses.

### Luminescence-based rates agree with CFU-based kill rates in 11 out of 22 antimicrobial assays

Given the linearity between light intensity and bacterial density shown above, we tested whether the rate of change of light intensity *ψ*_*I*_ aligns with the rate of change of bacterial population size (*ψ*_*B*_) under super-MIC antimicrobial concentrations. Since we cannot measure *ψ*_*B*_ directly, we first compared *ψ*_*I*_ to the CFU-based rate, *ψ*_CFU_, and then discussed their relation to *ψ*_*B*_. We measured CFU and light intensity over time for 20 drugs (Table S1) using an automated liquid handler (Methods) and estimated the distributions of the rates of change of CFU (*ψ*_CFU_) and light intensity (*ψ*_*I*_) by boot-strap (Figure 1, Table S2). We classified the luminescence and CFU-based rate distributions as “not significantly different” (n.s.) if each mean fell within the other’s 95% percentile and otherwise significantly different (*). All time-series data are presented in Figures S2–S20. For amoxicillin, cefuroxime, chloramphenicol, colistin, fosfomycin, penicillin, pexiganan, polymyxin B, rifampicin, and tetracycline, *ψ*_*I*_ and *ψ*_CFU_ did not differ significantly. However, we observed significant discrepancies for ampicillin, cefepime, ceftazidime, ciprofloxacin, doripenem, imipenem, mecillinam, meropenem, piperacillin, and trimethoprim.

**Figure 1.**
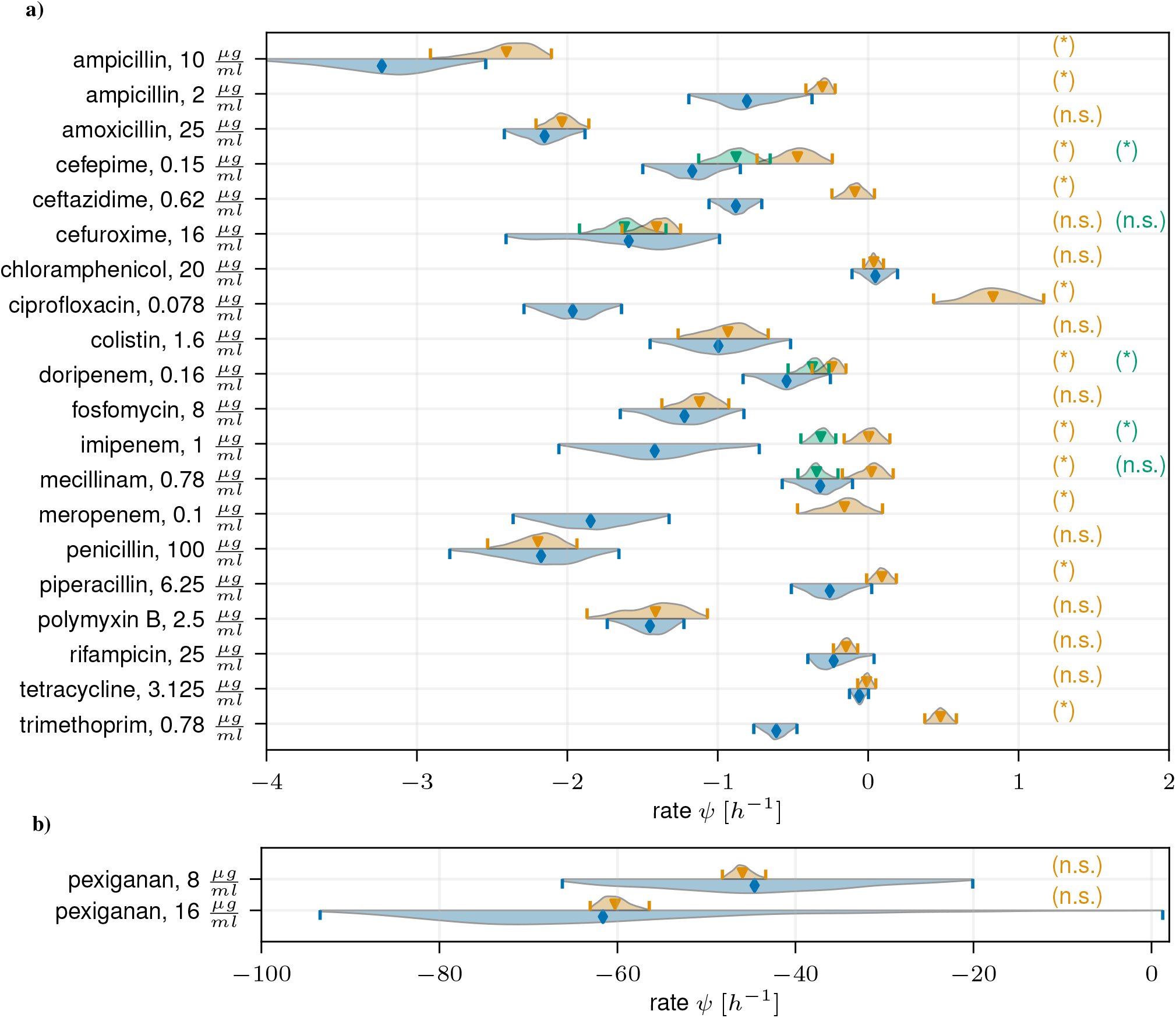
Comparison of CFU-based and luminescence-based rates of change. For each drug, we generated 2000 bootstrapped datasets by resampling time-series CFU and light-intensity data with replacement and fitted an exponential function to each bootstrap replicate to obtain distributions of rates. Panel (a) shows these distributions for 20 drug–concentration assays across 19 antibiotics; panel (b) shows the antimicrobial peptide pexiganan at 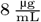 and 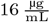. The distribution of *ψ*_CFU_ is shown in blue (lower half of each violin, diamond). The distribution of *ψ*_*I*_ is shown in orange (upper half, triangle). Green distributions (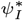; triangle) represent luminescence-based rates calculated from data starting at the first peak onward. Red distributions show volume-adjusted luminescence rates (*ψ*_*J*_; pentagon). Vertical lines mark the 95 % confidence intervals. Asterisk (*) or letters (n.s.) indicate whether CFU-based rates differ significantly from the corresponding luminescence-based rate or not (see Methods), with color coding matching the respective luminescence-based distribution. Wide confidence intervals for pexiganan reflect biphasic killing, steep curves, and noisy CFU data.

For all cases where significant discrepancies were observed, the light intensity declined more slowly than the CFU signal. In the following, we investigate potential reasons why the light signal may decline more slowly and the CFU signal more rapidly than the “true” *ψ*_*B*_.

### No support for SOS-driven increase in luminescence promoter activity

To explain the observed discrepancies between CFU- and luminescence-based rates, we tested whether the SOS response might upregulate lux expression, increasing the cell-specific luminosity. To this end, we exposed the strains to UV light to induce the SOS response. Specifically, we alternated between measuring light intensity and optical density, and exposing cultures to UV (Appendix S4).

UV treatment impaired bacterial growth significantly, yet the OD-normalized light intensity (∝ cell-specific luminosity) of UV-treated cells was lower than that of untreated controls (Figure S21d; t-test, *p* = 3 · 10^*−*5^ for the last time point). This implies that SOS induction did not upregulate lux expression, as cells under SOS had lower light output per cell than controls. While we cannot exclude the possibility that a non–UV-induced SOS response enhances cell-specific luminosity, our findings suggest that promoter upregulation is unlikely to explain why luminescence-based rates exceed CFU-based rates.

### Filamentation aligns with divergence between CFU- and luminescence-based rates

Antibiotic pressure is known to impair septation and induce bacterial filamentation. Our second hypothesis was that larger, filamented cells might emit more light per cell, thereby driving the divergence between CFU- and luminescence-based rates. We explored this for a subset of the antibiotics tested by imaging bacteria before and after two hours of antibiotic treatment (Figure S22–S34; see Methods). From these images, we measured bacterial length and width (Figure S35) and calculated the cell volumes (Figure 2, Appendix S4).

**Figure 2.**
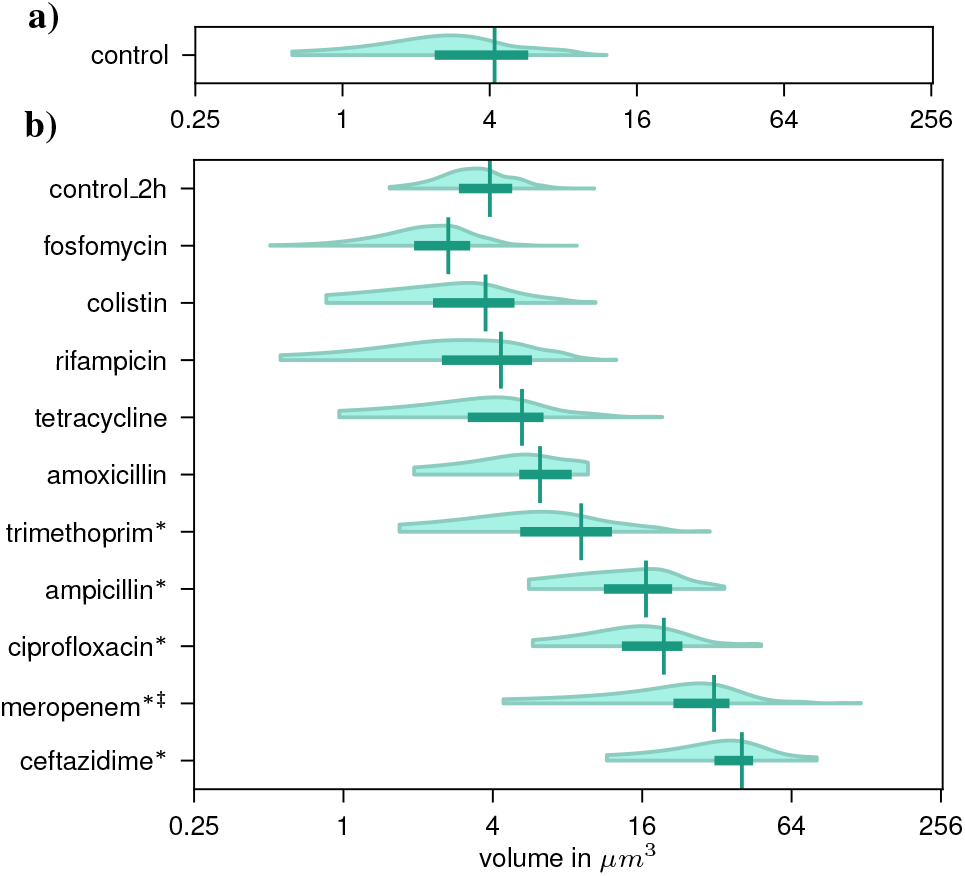
Density distributions (2.5–97.5% percentile range) of pooled cell volumes acquired by microscopy imaging, shown (a) before and (b) after 2 h of antibiotic treatment. Boxes indicate the 25–75 % interquartile range, and vertical bars mark the mean. Significance (*) was assessed by bootstrapping cell volumes 200 times with replacement for each replicate (image) and treatment, pooling the bootstrapped volumes by treatment, and comparing the resulting 95% confidence interval of the mean to that of the untreated control (`control_2h`). *‡* Carbapenems tend to deform cells into a lemon-like shape (Figure S30), resulting in poor fitting quality since our algorithm assumes a cylindrical geometry.

For antibiotics where microscopy showed no significant filamentation (amoxicillin, colistin, fosfomycin, rifampicin, and tetracycline, Table S3), luminescence-based and CFU-based rates did not differ significantly (Figure 1, Table S2). In contrast, for those drugs where microscopy data indicated significant filamentation (ampicillin, ceftazidime, ciprofloxacin, meropenem, and trimethoprim), luminescence-based rates were significantly higher than CFU-based rates.

### Filamentation model predicts divergence between luminescence- and CFU-based rates

To further investigate the link between filamentation and the recorded light signal, we developed a simplified population dynamical model which incorporates bacterial filamentation (Appendix S3). In this model, we assume that cell-specific luminosity scales with cell volume — i.e., the volume-specific luminosity remains constant. We simulate single-cell growth by assuming that biomass acquisition is independent of cell size, which leads to linear elongation [17]. A sudden reduction in the division rate by Δ*λ < λ*_0_ represents the onset of filament-inducing treatment and causes the mean cell volume to converge to a new, higher equilibrium (Figure S36).

As the mean cell volume stabilizes and cell-specific luminosity reaches equilibrium, the luminescence-based rate converges to the rate of change of population size *ψ*_*B*_ (Figure S36). Depending on the division and death rates, this dynamic can result in an initial peak in light intensity followed by a decline (Figure S36b). In the model, linear elongation is a mathematically convenient simplification. In reality, how cells elongate may depend on the specific strain, morphology, and treatment. However, as the initial peaks in light intensity arise from cell volume converging to a new equilibrium, non-linear elongation models can produce similar peaks.

We experimentally observed such peaks in light intensity for several drugs associated with filamentation, as predicted by the model for filamenting populations (Figures S3, S5, S9, S11, S12, and S13).

To investigate the dependence of *ψ*_*B*_ and *ψ*_*I*_ on treatment-induced changes in division rate and death rate (*δ*), we simulated four hours of treatment (details and parameters in Appendix S3). Our model shows that a reduction in *λ* leads to higher luminescence-based rates *ψ*_*I*_ relative to the rate of change of population size *ψ*_*B*_, with particularly large discrepancies when the death rate is low (Figure 3). The results from this model suggest that excluding early data points, where the mean cell volume changes rapidly, improves agreement between the estimated rates *ψ*_*I*_ and *ψ*_*B*_. This is evident in Figure S36b, where the slopes inferred from luminescence and from population size differ initially but are nearly identical at later times.

**Figure 3.**
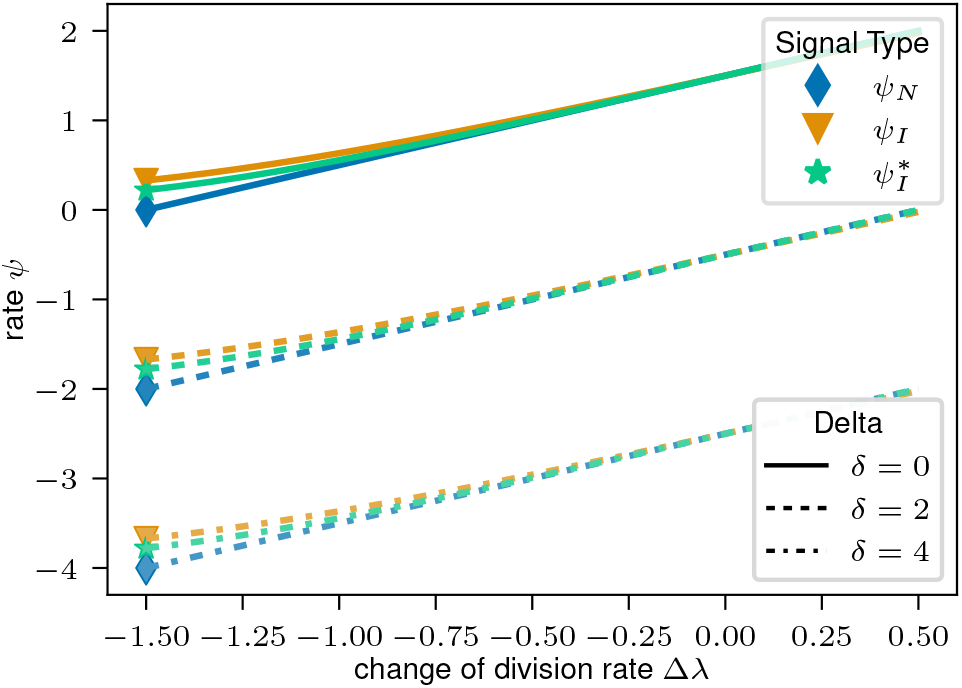
Simulations based on the filamentation model quantifying how changes in division rate due to treatment (Δ*λ*) and death rate (*δ*) influence the rate of change of population size (*ψ*_*B*_, blue), the rate of change of light intensity (*ψ*_*I*_, orange), and the rate of change of light intensity when the first 2 hours of data are excluded (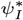, green). These illustrative simulations were conducted using an initial division rate *λ*_0_ = 1.5 h^*−*1^. More details and all parameter values can be found in Appendix S3.

We tested this approach by refitting all experimentally acquired luminescence-based rates that exhibited an initial peak, excluding data points recorded before the light signal reached its maximum. The resulting distributions of 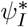 (green) are shown in Figure 1. An exception was made for meropenem, for which we know the change of cell volume and thus applied an alternative correction as described below. This adjustment substantially reduced the difference between CFU- and luminescence-based estimates for all tested drugs and fully eliminated the discrepancy for mecillinam.

### Adjusting luminescence intensities by changes in volume narrows the gap between CFU- and luminescence-based rates

Given the model-predicted differences between *ψ*_*I*_ and *ψ*_*B*_ in filamenting populations, we next tested whether combining morphological data with measured light intensities can help infer *ψ*_*B*_. This approach only works if the volume-specific luminosity is constant (Appendix S2, Equation S12). We used the mean cell volume data acquired by microscopy imaging before (*v*_obs,0_, Figure 2a), and after treatment (*v*_obs,2h_, Figure 2b), for all drugs that caused significant filamentation (ampicillin, ceftazidime, ciprofloxacin, meropenem, and trimethoprim). The light intensities were then volume-corrected as *J* (*t*) = *I*(*t*) *v*_obs,0_*/v*(*t*), where *v*(*t*) is derived from the filamentation model (Eq. (S44), Appendix S3). The free parameters were determined by minimizing Equation S51. All adjusted light signals *J* are shown in Figures S2a, S4, S7, and S13.

For ampicillin, the CFU-based and volume-corrected luminescence-based rates (*ψ*_*J*_) did not differ significantly (Table S2). For ceftazidime, ciprofloxacin, and meropenem, volume correction reduced the discrepancy, but *ψ*_*J*_ remained significantly above *ψ*_CFU_. We observed in all experiments that volume correction narrowed but never reversed the discrepancy (*ψ*_CFU_ ≤ *ψ*_*J*_ ≤ *ψ*_*I*_, Equation S23). From this observation and the derivation in the SI (Appendix S2), we conclude that *ψ*_*I*_ is closer to *ψ*_*V*_ (rate of total cell volume change) than to *ψ*_*B*_ (rate of bacterial number change).

Three factors may explain these residual differences: (i) the assumption of constant volume-specific luminosity may not hold; (ii) the approximation of *v*(*t*) may be inaccurate, and excluding entangled or overlapping cells from the analysis introduces a bias that underestimates the volume of heavily filamented cells; (iii) the CFU-based method may underestimate *ψ*_*B*_.

For ciprofloxacin, the large disparity between *ψ*_CFU_ and *ψ*_*I*_ is unlikely to be explained by factors (i) and (ii) alone. Bringing the two rates into agreement solely by adjusting cell-specific luminosity would require an almost four-order-of-magnitude increase, which appears implausible. We therefore conclude that, in this case, CFU-based measurements likely overestimate the rate of population decline, corresponding to an underestimation of *ψ*_*B*_ (iii).

### CFU-based estimates can overestimate the rate of population decline

After exploring why luminescence assays can underestimate the rate of population decline, we now explore why CFU assays may overestimate it. The rate of change of the CFU signal, *ψ*_CFU_, only matches *ψ*_*B*_ if the number of colonies emerging per plated bacterium, *η* ∈ [0, 1] (Equation S4), is constant over time (Appendix S2, Equation S6). This assumption can be problematic for three main reasons:

#### a) Loss of culturability

The number of colonies emerging per plated bacterium, *η*, depends on the division rate *λ*, which can be affected temporarily or permanently by treatment ([18]), e.g. due to DNA damage. In extreme cases, viable and culturable cells can be converted into viable but non-culturable (VBNC) cells ([19–21]), meaning they continue to be metabolically active but cease to divide (*λ* = 0) and therefore no longer form colonies on agar. Typically, cells can reproduce at the start of a time-kill assay but may, depending on the drug, partially or completely lose this ability as the assay progresses, causing an underestimation of *ψ*_*B*_.

#### b) Antimicrobial carryover

Antibiotics transferred onto agar by plating a diluted culture can have a residual treatment effect on either the division rate *λ* or the death rate *δ*, depending on the mode of action of the drug, thereby reducing the probability of colony formation. This phenomenon, known as antimicrobial carryover, has been described in previous studies [22–24]. Its effect is usually minimal at the start of a time-kill assay, when bacterial density is high and plated samples are highly diluted. However, as the assay progresses and bacterial density declines, less dilution is needed, increasing the concentration of the transferred antibiotic. As a consequence, CFU-based rates would underestimate *ψ*_*B*_ between two time points *t*_0_ and *t*_1_ by 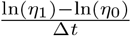, where *η*_*i*_ denotes the mean number of colonies formed per plated bacterium at time *t*_*i*_.

#### c) Aggregation

Filamentation or altered cell adhesiveness can change the distribution of colony-initiating cluster sizes on agar after plating (Equation S3). Changes in cluster size in turn, affect the average number of clusters per plated bacterium, thereby biasing estimates of *ψ*_*B*_.

### Partial loss of culturability causes CFU to underestimate *ψ*_*B*_ for ciprofloxacin and trimethoprim treatment

During ciprofloxacin treatment, CFU counts fell steeply while light intensity continued to rise (Figure S7). This discrepancy is consistent with previous reports comparing CFU and luminescence during fluoroquinolone killing [14, 25]. To investigate the cause of this discrepancy, we plated treated cultures on phosphate-buffered saline (PBS) agar containing propidium iodide (PI), a red fluorescent dye that binds to nucleic acids but cannot penetrate intact cell membranes. Microscopic imaging (Figure S27) revealed almost no red fluorescence, indicating that the cells remained impermeable. Although impermeability alone does not confirm viability, additional observations support the conclusion that most cells were still alive: the absence of bacterial debris (as has been observed for drugs with similar decline in CFU such as amoxicillin), visible growth indicated by increased cell size compared to two hours earlier, and continued (and even increased) light emission. These findings suggest that most cells remain alive but are unable to form colonies under the provided conditions, possibly due to DNA damage induced by ciprofloxacin [26]. This observation aligns well with previous studies on ciprofloxacin, which found that CFU can underestimate viability relative to non-culture-based methods [19, 27, 28].

Trimethoprim treatment showed similar, though less pronounced, results (Figure S19). Trimethoprim, which impairs DNA replication ([29]), likewise caused an increase in light intensity and a decline in CFU counts, while microscopy revealed intact, mostly impermeable, filamented cells (Figure S34).

### Antimicrobial carryover causes underestimation of *ψ*_*B*_ for pexiganan using CFU

Building on our understanding of when luminescence assays accurately estimate *ψ*_*B*_, we hypothesized that antimicrobial peptides (AMPs) would be an ideal application for this method. We expected that, during the AMP’s short killing phase, changes in the cell-specific luminosity would remain negligible compared to the high kill rates AMPs can achieve.

Initially, however, we failed to recover almost any colonies on agar, despite the light intensity indicating a high enough bacterial density. Moreover, colony counts were inconsistent across dilutions: 100-fold and 1000-fold dilutions from the same cultures yielded similar colony numbers, instead of reflecting the tenfold difference. We suspected that AMPs from the liquid culture, including those attached to the bacterial surface, were carried over into the PBS dilution medium, causing continued cell death during dilution and after plating. To test this, cultures treated with pexiganan for 1 min were diluted 1:100 in PBS supplemented with various concentrations of CaCl_2_ and MgCl_2_. These compounds were selected based on prior evidence that they inhibit the activity of other AMPs ([30]). We sampled and plated at four time points from these diluted cultures, approximately 45 minutes apart.

Our results show that supplementing the dilution medium increased the measured CFU substantially (Figure S38, Appendix S4). Conversely, diluting in unsupplemented PBS did not stop bacteria from dying. Supplementing 100 mM MgCl_2_ yielded the highest CFU count for the first time point (Table S4). Since CFU cannot systematically overestimate bacterial density, this count represents the best estimate of the bacterial density. Consequently, we supplemented the PBS with 100 mM MgCl_2_ in subsequent pexiganan experiments. Given this insight into the residual killing effect of pexiganan and how to mitigate it, we repeated the CFU time-kill experiment using two different dilution media: pure PBS and PBS supplemented with 100 mM MgCl_2_ (Appendix S4). We recorded three replicates for each of the two time-kill curves and counted colonies on all agar plates from three dilution steps for both dilution media. To lower the detection limit by one order of magnitude, we increased the plated volume from 10 µL to 100 µL. Since this volume exceeds the capacity of the automated high-throughput setup, we used the standard manual CFU plating method instead.

We observed a much steeper initial decline in CFU for the cultures diluted in pure PBS compared to the supplemented ones (Figure S39). In pure PBS, more highly diluted samples consistently yielded higher CFU estimates (Figure S39a), supporting the antimicrobial carryover hypothesis. This pattern diminished over time, suggesting a reduction in the residual killing effect of pexiganan, which we discuss below.

### Luminescence and CFU show identical decline rates for pexiganan time-kill curves if residual killing is prevented

To confirm that eliminating residual pexiganan killing aligns CFU and luminescence, we supplemented PBS with 100 mM MgCl_2_ and measured both signals at pexiganan concentrations of 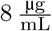 and 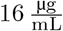 using the “rapid luminescence-CFU assay setup” (Methods). We observed no significant difference between the CFU- and luminescence-based rates for either of the tested pexiganan concentrations (Figure 1b, Table S2). However, examining the time series (Figure S20) revealed that while CFU and luminescence signals declined in parallel for the 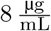 treatment, they diverged for the 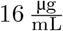 kill curve. In this case, the CFU signal initially declined much faster (rates below *−* 200 h^*−*1^), and subsequently declined more slowly than the corresponding luminescence signal, ultimately resulting in a similar average rate (approximately *−*60 h^*−*1^).

### Pexiganan rapidly loses killing capacity

During the pexiganan experiments, we observed an initial steep decline in bacterial density, predicted by both CFU and luminescence, followed by almost constant signals (Figures S20, S39). Two possible, non-exclusive explanations for this observation are: first, pexiganan is deactivated or sequestered from the medium by attaching to targets on the bacteria over time; and second, the remaining bacteria are unaffected by the AMP because they are resistant or persisters. To investigate the first explanation, we exposed bacteria to pexiganan for 5 min at 16^µg^, after which the supernatant was collected and tested for its ability to kill bacteria (Appendix S4). While the initial treatment showed rapid bacterial killing (*ψ*_CFU_ ≈ 46 *−* h^*−*1^ between *t*_0_ = 0 and *t*_1_ = 5 min), bacteria exposed to the supernatant alone showed no significant reduction in viability (Figure S40, Table S5). These results show that the supernatant has no residual killing effect. This makes deactivation or sequestration through the attachment of pexiganan the likely explanation for the flattening CFU signal, even though we did not assess whether the surviving cells are resistant or persisters.

## Discussion

We evaluated whether luminescence can serve as a high-throughput proxy for population dynamics by comparing it with CFU assays. We found no significant difference between CFU- and luminescence-based rates for treatments that neither induce substantial changes in culturability nor provoke strong morphological changes, such as filamentation.

However, for drugs that induce filamentation and/or loss of culturability, the two methods can yield significantly different results. The divergence between the two rates does not imply that either method is incorrect; rather, CFU and luminescence capture different population properties.

The CFU method counts bacteria capable of forming colonies on permissive media (i.e., culturable cells). When inferring growth rates from CFU, the observed rate of change *ψ*_CFU_ reflects both the rate of change of population size *ψ*_*B*_ and changes in the probability that a plated bacterium forms a colony, *η* (Equation S4). The CFU-based rate equals *ψ*_*B*_ only if *η* remains constant over time.

However, *η* can change for three main reasons. First, altered clustering behavior can change how many bacteria seed a single colony. This directly shifts the observed colony count. Second, physiological changes to the bacteria may lead to a temporary or permanent change in culturability ([18–21, 27, 31]), which may increase the fraction of viable cells that fail to form colonies, e.g. due to a reduced division rate. Third, residual drug activity carried over to the agar can alter on-plate conditions, thereby reducing division or increasing the death rate ([22–24]).

Preventing antibiotic carryover when handling low-density cultures treated with highly concentrated antimicrobials is challenging and, in some cases, infeasible. Centrifugation-based washing (pelleting bacteria and replacing the supernatant) can remove residual drug, but only if the processing delay is negligible relative to the antibiotic’s killing kinetics — a condition unlikely to hold for fast-acting agents such as AMPs. Moreover, although bacteria generally tolerate high centrifugal forces ([32]), the impact on compromised cells, such as those with destabilized walls, is unknown. As an alternative, we found that supplementing the dilution medium with MgCl_2_ effectively neutralizes residual AMP activity, preventing residual killing effects in CFU assays for pexiganan. However, this strategy is not generalizable, as for many antimicrobials the corresponding deactivating agents are unknown — or may not even exist. For these cases, it may be impossible to accurately derive *ψ*_*B*_ from CFU counts at high drug concentrations.

In contrast to CFU assays, luminescence assays become more reliable at high, fast-killing concentrations. This is because *ψ*_*I*_ reflects both the rate of population size change, *ψ*_*B*_, and changes in cell-specific luminosity (Equation S9); when population declines rapidly, *ψ*_*I*_ converges to *ψ*_*B*_.

One potential exception is the delay between the decline of CFU and the decline of the luminescence signal, observed during the pexiganan experiments. This discrepancy may arise from two effects: either the CFU signal declines more steeply, or the luminescence signal declines more slowly than the number of living cells. CFU may overestimate the decline, as damaged but living cells have a reduced probability of forming colonies. Conversely, luminescence may under-estimate the decline if there is a delay between cell death and cessation of luminescence. For antimicrobials that lyse cells (such as pexiganan), we would expect a rapid, though not instant, cessation of luminescence due to the dilution of all reactants. The delay may be more pronounced for drugs that kill without lysing cells. However, metabolically active and impermeable cells are difficult to characterize as dead in the first place. Based on our experimental data, we cannot distinguish between these possibilities and therefore cannot exclude that luminescence assays underestimate extremely rapid kill rates.

For lower kill rates, we observed that the absence of filamentation was a good indicator of stable cell-specific luminosity. In cases where drugs induced filamentation, CFU- and luminescence-based rates diverged. We further demonstrated that correcting for the increased cell size partly compensates for the difference between CFU- and luminescence-based estimates, and we found that the rate of change in light intensity is closer to the change in total cell volume than to the change in total cell number (Appendix S2). This makes a constant volume-specific (or mass-specific) luminosity a better assumption than a constant cell-specific luminosity.

Changes in both CFU and luminescence are used as proxy signals for population growth rates ([1, 2, 8–10, 12, 13]). Whether discrepancies between the changes in these proxy signals and the changes in living bacteria pose a problem depends on the underlying biological question. The rate of change of living bacteria, *ψ*_*B*_, is most commonly applied in theoretical modeling to create predictions, making its estimation important. If, instead, the aim is to assess a population’s reproductive potential, for example, in studies focusing on evolutionary dynamics, examining changes in the number of culturable cells (as approximated by CFU) may be more relevant than *ψ*_*B*_, as only culturable cells contribute to subsequent generations and thus to evolution. Tracking changes in total biomass (closer to *ψ*_*I*_) can be more relevant than the number of living bacteria, as biomass accounts for a potential “catch-up” effect, whereby filamented cells fragment into multiple viable units once antibiotic pressure is removed ([31, 33]).

A notable challenge of using luminescence assays is the absence of a fundamental biological principle linking light intensity uniquely to a single population property. Beyond cell number and biomass, the light intensity can depend on treatment-induced metabolic changes, which potentially explain some of the discrepancies between luminescence- and CFU-based rates observed. Furthermore, the availability of nutrients can influence luminosity, limiting assays to time-frames during which the nutritional availability remains stable.

A further subtlety arises from within-population heterogeneity: cell-specific luminosity can vary between individual bacteria. Such variation may bias population-level estimates if it co-varies with the susceptibility to antimicrobials.

Measuring population decline is challenging. In this work, we addressed some of the complexities involving the luminescence method, but several questions remain open for follow-up work. Our study was motivated by the question of whether changes in light intensity follow those in cell number, and we therefore used CFU as the reference metric. Supplementing these experiments with microscopy, we found that light intensity tracks cumulative cell volume more closely than cell number. A natural next question is therefore how closely luminescence tracks biomass — addressing it, however, requires a reference method designed to capture volume or mass directly, rather than cell count.

Generalizability beyond *E. coli* is a second open question: while the core principle that larger cells emit more light should hold broadly, the morphological and physiological responses to antimicrobial treatment may be strain-specific, so the drug-specific observations do not necessarily transfer across species. A related caveat is that most drugs in this study were tested at a single concentration, leaving the concentration dependence of these drug-specific findings still to be established.

Our results show that neither CFU nor luminescence is optimal for every experimental scenario. Instead, CFU and luminescence work best under different conditions, measure different population properties, and complement each other. At low and intermediate drug concentrations, changes in CFU accurately reflect changes in bacterial density, but CFU becomes unreliable at high drug concentrations.

Luminescence, by contrast, becomes more reliable at high concentrations, where CFU becomes unreliable. In practice, the luminescence method significantly reduces labor, consumables, and costs: eight PD curves with twelve concentrations and four replicates each can be fit on a single 384-well plate, whereas measuring CFU would require more than 8,000 agar plates, hundreds of dilution plates, and substantial manual labor. Given their scalability and cost-effectiveness, luminescence assays offer a valuable alternative for high-throughput analysis, particularly at high antimicrobial concentrations, where traditional methods become unreliable or even unusable.

## Methods

### Strains

We generated a bioluminescent strain by integrating a modified *P. luminescens luxCDABE* operon, driven by the constitutive *λ*-Pr promoter, together with a kanamycin resistance cassette (as a marker) into the chromosome of *Escherichia coli* MG1655. This integration replaced the galK gene and was achieved using *λ*-Red recombination ([34]), following a protocol by Hughes [35, 19–26]. The integrated elements were derived from the pCS-*λ* plasmid ([8, 36]). Primers are listed in Table S6. For all time-course experiments, we prepared three replicate exponential cultures by diluting overnight cultures (grown for approximately 18 hours) 1:100 and growing them to exponential phase for 1–1.5 hours.

### Media

We used LB (Sigma L3022) as a liquid medium and, as a solid medium for CFU plating, LB with 1.5 % agar. Cultures were treated by diluting the tenfold working concentration of one of 20 antimicrobials 1:10. All MICs, determined by broth microdilution ([37]), and the concentrations used are listed in Table S1. Working concentrations were centered around the MIC, with some variation due to rounding convenience and variability in repeated MIC tests. For colistin and polymyxin B, we used lower concentrations, as higher concentrations in our setup consistently yielded too few colonies for meaningful analysis.

Phosphate-buffered saline (PBS, Sigma 79383) was used as the diluent for CFU assays. If cultures were treated with pexiganan, 100 mM MgCl_2_ was added. For microscopy, we added 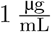 propidium iodide (PI) to the liquid medium and used PBS/PI agar plates (containing PBS with 1.5 % agar and 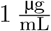 PI) as solid medium.

### Automated CFU plating

We automated the high-throughput colony-count method described by [38] using an Evo 200 liquid-handling platform (Tecan) integrated with a Liconic STX100 incubator. The platform handles liquids and automatically moves, images, and incubates plates. We produced six colony streaks by spotting six 10 µL drops of diluted bacterial culture onto a one-well agar plate. Plates were automatically tilted for 7 s on a custom-built tilter integrated into the platform, to spread the drops and distribute the bacteria. After incubation, plate images were captured using the Pickolo camera (SciRobotics).

The platform is controlled by custom-generated worklists executed in the native software “Evoware”. These worklists were generated using the Python package `pypetting` (version 1.0.1). We analyzed the captured images of the agar plates using a custom colony-recognition script that automatically identifies colonies and allows the manual addition of unidentified colonies and the removal of mismatched ones. All Python classes for generating the worklists and analyzing colonies are available at Zenodo (DOI: 10.5281/zenodo.15261184).

### Luminescence measurements

To record the luminescent light intensity, we used an Infinite F200 spectrophotometer plate reader (Tecan), which is also integrated into the liquid-handling platform, with an exposure time of one second. We set 20 rlu as the lower detection limit and excluded all data points below.

### Luminescence-CFU assay setup

To measure the CFU and light intensity at seven time points, we treated the exponential cultures and then distributed them onto seven (one for each time point) white 384-well plates (Greiner, 781073), with each culture well containing 54 µL medium and 6 µL 10x stock solution. We adjusted the duration of the experiments between two and five hours, depending on the anticipated kill rate. For each time point, an assay plate was transferred from the incubator to the plate reader for luminescence measurement. Subsequently, a dilution series was conducted directly in the white plate and plated using the automated plating method, after which the plate was discarded.

### Rapid luminescence-CFU assay setup

This experiment is a variation of the *Luminescence-CFU assay setup*, adjusted to measure rapid kill curves for the antimicrobial peptide pexiganan. In this setup, we captured four time points within 5 min. Cultures were treated in a 96-deep-well plate (Greiner, 780285) by adding 100 µL of the 10x stock solution to 900 µL exponential phase culture. 60 µL of the treated culture was then transferred to a 384-well white plate (Greiner, 781073) and placed in the plate reader for continuous luminescence recording. For the four CFU time points, samples were taken directly from the deep-well plate, automatically diluted in PBS supplemented with 100 mM MgCl_2_ in a 96-well plate (Greiner, 655101) to halt the antimicrobial activity and then plated.

### Morphology experiments

To assess treatment-induced morphological changes, we imaged treated (for 2 hours) and untreated bacteria by spotting 2 µL droplets onto PBS/PI-agar plates. The spots were cut out and flipped onto Ibidi *µ*-dishes (Ibidi, 80136) for imaging. We used an Eclipse Ti2 microscope (Nikon) with a 100x objective connected to a DS-Qi2 Nikon Scientific CMOS (sCMOS) camera to image the bacterial cells. The microscope setup included an additional 1.5x zoom, which was used only for some images due to unintentional variation. We estimated the width and length of the bacterial cells using a custom Python script, as described in Appendix S4.

### Fitting rates of change *ψ*

To compare the rates of change of two signals, we first excluded all data below the detection limits (empty plates or light intensity below 20 rlu). We then truncated both signals at the latest time point where both remained above the detection limit, ensuring the same time frame was used for comparison. Next, we bootstrapped 200 datasets with replacement per signal, while ensuring that each dataset contained more than one time point. For each dataset, we applied a simple regression to fit an exponential function to all time points of each time-kill curve, resulting in distributions with 200 rate estimates each.

### Significance

We classify two distributions of rates as not significantly different (n.s.) if the mean of each distribution falls within the 95% confidence interval of the other. Otherwise, we classify them as significantly different (*).

## Data, Materials, and Software Availability

Experimental datasets are available at Zenodo (DOI: 10.5281/zenodo.15261454). Analysis scripts, plate-handling worklists, colony-recognition code, and model code are available at Zenodo (DOI: 10.5281/zenodo.15261184).

## Acknowledgements

We thank Marco La Fortezza and Ricardo León Sampedro for assistance with microscopy imaging. During manuscript preparation, we used OpenAI’s ChatGPT for editorial assistance (grammar, phrasing, and proofreading). This work was supported by funding from ETH Zurich.

