## Supplementary material for "High-Throughput Quantification of Population Dynamics using Luminescence"

### S1 Nomenclature

| | |
|---|---|
| $B$ | number of living bacteria in a well |
| $v$ | cell-specific mean volume |
| $V$ | cumulative cell volume, $V = vB$ |
| $I$ | measured light intensity (detector-recorded bioluminescence signal) |
| $J$ | volume-corrected bioluminescence signal, $J = I\, v_0/v$ |
| $\kappa$ | instrument-specific transfer factor (fraction of emitted light detected) |
| $\theta$ | volume-specific luminosity (bioluminescence per cell volume) |
| $\phi$ | cell-specific luminosity (bioluminescence per cell) |
| $\omega$ | OD-normalized light intensity ($I$ divided by measured optical density) |
| $\lambda$ | bacterial division rate |
| $\gamma$ | biovolume acquisition rate |
| $\delta$ | bacterial death rate |
| $\tau$ | treatment effect |
| $b$ | bacterial density |
| $c$ | drug concentration |
| $p_E$ | extinction probability of a lineage originating from a single plated bacterium |
| $p_C$ | survival probability of a lineage from a single plated bacterium ($p_C = 1 - p_E$) |
| $\eta$ | number of colonies per plated bacterium ($\eta \in [0,1]$) |
| $V_w$ | well volume |
| $\epsilon$ | volume increment |
| $\psi_X$ | rate of change of population property $X$ ($\psi_X = d\ln(X)/dt$), where $X \in \{B, V, I, J, \mathrm{CFU}\}$ |

### S2 Mathematical descriptions

#### S2 a Light-related terminology.

In this manuscript, *total luminosity* ($L$) refers to the total light output produced by $B$ bacteria of a *bioluminescent* bacterial culture in a well with volume $V_w$. During the *luminescence assays*, we capture a fraction $\kappa$ of the total luminosity ($\phi B$) as light intensity $I(t) = \kappa\phi B$. We call the exponential decline rates based on these intensities *luminescence-based rates*, $\psi_I$. We use *cell-specific luminosity* ($\phi$) for light output per cell and *volume-specific luminosity* ($\theta$) for light output per unit cell volume. When $I(t)$ is normalized by the measured optical density, we obtain $\omega$, the *OD-normalized light intensity*. $J(t)$ is the light intensity $I(t)$ adjusted by the relative change in cell volume $\frac{v_0}{v(t)}$.

#### S2 b Testing for linearity between bacterial density and luminescent light intensity.

To test whether bacterial density and bacterial luminescent light intensity are linearly related, we grew three replicate overnight cultures. To replenish nutrients, we diluted each culture 1:10 in fresh LB medium and incubated for 60 minutes. We subsequently performed a 10-fold dilution series in a 384-well white plate (Greiner, 781073) and immediately measured the light intensity (see Fig. 1 in the main text). Each plate included wells containing only medium to determine a blank (median $\approx 37.33\,\mathrm{rlu}$), which we subtracted from all measurements.

We assessed linearity between light intensity ($I$) and bacterial density ($b$) by fitting a linear model without intercept on the original scale,

$$I_i = m \cdot b_i,$$

which corresponds to a log–log regression with intercept:

$$\ln(I_i) = \ln(m) + \ln(b_i).$$

Summing over all $n$ observations gives

$$n\ln(m) = \sum_{i=1}^{n} \ln(I_i) - \sum_{i=1}^{n} \ln(b_i),$$

The optimal conversion factor $m$ is then

$$m = \frac{\left(\prod_{i=1}^{n} I_i\right)^{\frac{1}{n}}}{\left(\prod_{i=1}^{n} b_i\right)^{\frac{1}{n}}},$$

yielding $m = 0.006\ \frac{\mathrm{rlu\,ml}}{\mathrm{CFU}}$.

We observed a linear relationship ($R^2 = 0.987,\ F = 1164.14,\ p < 10^{-14}$) between bacterial density and light intensity for intensities above $20\,\mathrm{rlu}$ ($\sim 3 \times 10^2$ CFU/ml). Consequently, we use $20\,\mathrm{rlu}$ as the lower detection limit for luminescence. Additionally, we conclude that $\kappa$ is independent of the bacterial density (no overshadowing effects), within the relevant range of densities. Since the plate reader setup remains constant within an experiment, we assume $\kappa$ to be constant for following analyses.

#### S2 c Rate of change of CFU count.

To infer the rate of change of CFU, we assume bacteria spend only a short time in the low-nutrient dilution medium, so replication and death are negligible during that phase.

Most colonies originate from clusters only comprising a single cell, but some from founding clusters of multiple cells. Let $f(n)$ be the probability that a randomly chosen cluster contains $n$ bacteria. A cluster of size $n$ forms a colony with probability

$$1 - (p_E)^n, \tag{S1}$$

where $p_E$ is the probability that a lineage originating from a single bacterium goes extinct (extinction probability). The mean probability that a plated cluster forms a colony is:

$$\bar{g} = \sum_{n=1}^{N} f(n)\,(1 - (p_E)^n). \tag{S2}$$

The mean cluster size is

$$\bar{n} = \sum_{n=1}^{N} n\,f(n). \tag{S3}$$

The mean number of colonies emerging per plated bacterium can be approximated by:

$$\eta = \frac{\bar{g}}{\bar{n}}, \qquad \eta \in [0,1]. \tag{S4}$$

The predicted CFU per ml given a bacterial density $b = \frac{B}{V_w}$, with $B$ the number of bacteria per well and $V_w$ the constant well volume, is:

$$\mathrm{CFU} = \eta \frac{B}{V_w}. \tag{S5}$$

Taking the logarithmic derivative yields:

$$\psi_{\mathrm{CFU}} = \frac{d}{dt}\ln(\mathrm{CFU}) = \frac{d}{dt}\ln(\eta) + \frac{d}{dt}\ln(B). \tag{S6}$$

In practice $\eta$ is often unknown. We can only estimate $\psi_B$ from CFU data if we assume that $\eta$ is constant over time.

$$\psi_{\mathrm{CFU}} = \frac{d}{dt}\ln(B) = \psi_B. \tag{S7}$$

To justify a constant $\eta$ we must assume that the cluster-size distribution $f(n)$ and the extinction probability $p_E$ do not change over time. Both assumptions may fail, for example if cells filament or if the division or death rate change. We discuss the behaviour of $p_E$ below.

#### S2 d Rate of change of luminescence.

The observed light intensity $I$ is a fraction $\kappa$ of the total luminosity ($\phi B$) of a bioluminescent culture, where $\phi$ is the cell-specific luminosity (the amount of light emitted by one bacterium). The light intensity can thus be written as:

$$I = \kappa \phi B. \tag{S8}$$

The rate of change of light intensity is therefore:

$$\psi_I = \frac{d}{dt}\ln(I) = \frac{d(\ln(\kappa) + \ln(\phi) + \ln(B))}{dt}. \tag{S9}$$

We assume that $\kappa$ remains constant over time, as we explained above ("Testing for linearity between bacterial density and luminescent light intensity"). If we assume that the cell-specific luminosity $\phi$ is also constant, $\psi_I$ equals the rate of bacterial count change $\psi_B$, as Equation S9 simplifies to:

$$\psi_I = \frac{d}{dt}\ln(I) = \frac{d}{dt}\ln(B) = \psi_B. \tag{S10}$$

#### S2 e Rate of change of volume-corrected luminescence.

Alternatively, we can link the measured light intensity to the number of bacteria using the mean cell-specific volume ($v$) and the volume-specific luminosity ($\theta$):

$$I = \kappa \theta v B. \tag{S11}$$

Defining the volume-corrected luminescence as $J(t) = I(t)v_0/v(t)$, we can compute its rate of change as:

$$\psi_J = \frac{d(\ln(I \cdot v_0/v))}{dt} = \frac{d(\ln(\kappa) + \ln(v_0) + \ln(\theta) + \ln(B))}{dt}. \tag{S12}$$

If we assume that the volume-specific luminosity $\theta$ is constant, this estimate equals the rate of change of the number of living bacteria ($B$):

$$\psi_J = \frac{d}{dt}\ln(I \cdot v_0/v) = \frac{d}{dt}\ln(B) = \psi_B. \tag{S13}$$

#### S2 f Change of light intensity is closer to the rate of change of total cell-volume than to the rate of change of number of bacteria.

We made two empirical observations under all tested drug conditions:
**(i)** *For the subset of drugs imaged using microscopy, the mean specific cell volume never significantly decreased between the first and second time point*:

$$\frac{d\ln v}{dt} \geq 0 \tag{S14}$$

The rate of change of total cell volume is given by:

$$\psi_V = \frac{d\ln(vB)}{dt} = \frac{d\ln v}{dt} + \psi_B. \tag{S15}$$

From this relation and observation **(i)**, we directly obtain:

$$\boxed{\psi_V \geq \psi_B} \tag{S16}$$

**(ii)** *The rate of change of volume-corrected light intensity was never significantly lower than the corresponding rate of change of CFU*:

$$\psi_J = \frac{d\ln J}{dt} \geq \psi_{\text{CFU}} \tag{S17}$$

Since we can express the light intensity as $I = \kappa\theta v B$, the volume-corrected luminescence rate becomes:

$$\psi_J = \frac{d}{dt}\ln(\kappa\theta v_0 B) = \psi_B + \frac{d\ln\theta}{dt}. \tag{S18}$$

Observation **(ii)** thus implies:

$$\psi_B + \frac{d\ln\theta}{dt} \geq \psi_{\text{CFU}} \tag{S19}$$

If we make the assumption that the rate of change of CFU equals that of bacterial count ($\psi_{\text{CFU}} = \psi_B$) we obtain:

$$\frac{d\ln\theta}{dt} \geq 0 \tag{S20}$$

Using the rate of change of light intensity we get:

$$\psi_I = \frac{d\ln(\kappa\theta v B)}{dt} = \psi_V + \frac{d\ln\theta}{dt}. \tag{S21}$$

From this relation and Equation S20, we follow:

$$\boxed{\psi_I \geq \psi_V} \tag{S22}$$

Combining Equation S16 and Equation S22 yields:

$$\boxed{\psi_I \geq \psi_V \geq \psi_B} \tag{S23}$$

allowing us to conclude that, during our experiments, the luminescence-based rate is closer to the rate of change of total cell volume (likely identical to the rate of change of biomass) than to the rate of change of bacterial count.

#### S2 g Colony formation – birth-death Markov Model.

We use a basic birth-death Markov model, as described by [1], [2], [3], and [4], to model the probability that a single plated bacterium creates a colony.
In this model, as in [1], $P_0(t)$ describes the probability that the population originating from this single bacterium goes extinct by the time $t$:

$$P_0(t) = \frac{\delta}{\lambda} \cdot \frac{E(t) - 1}{E(t) - \frac{\delta}{\lambda}}, \tag{S24}$$

where $\delta$ is the death rate, $\lambda$ the division rate, and $E(t) = e^{(\lambda-\delta)t}$. For $t \to \infty$, $P_0(t)$ converges to the extinction probability $p_E$ for a lineage originating from a single cell.

$$p_E = \begin{cases} 1, & \text{if } \lambda \leq \delta, \\ \frac{\delta}{\lambda}, & \text{if } \lambda > \delta. \end{cases} \tag{S25}$$

#### S2 h Colony formation for bacteriostatic and bactericidal drugs.

In the following, we call the death and division rate in the absence of treatment $\delta_0$ and $\lambda_0$, respectively, and the treatment-induced increase in death and reduction in division rate $\delta_T$ and $\lambda_T$, respectively. We then rewrite the net growth rate as:

$$\psi = \lambda_0 - \lambda_T - \delta_0 - \delta_T \tag{S26}$$

Furthermore, we define the combined treatment effect:

$$\tau = \psi_0 - \psi = \delta_T + \lambda_T \tag{S27}$$

We write the probability of colony formation (from a single cell) as:

$$p_E = \begin{cases} 1, & \text{if } \lambda_0 - \lambda_T \leq \delta_0 + \delta_T, \\ \frac{\delta_0+\delta_T}{\lambda_0-\lambda_T}, & \text{if } \lambda_0 - \lambda_T > \delta_0 + \delta_T. \end{cases} \tag{S28}$$

We then define the extinction probability for purely bacteriostatic drugs ($\delta_T = 0$ and $\lambda_T = \tau$) as:

$$p_{E,\text{stat}} = \begin{cases} 1, & \text{if } \lambda_0 - \tau \leq \delta_0, \\ \frac{\delta_0}{\lambda_0-\tau}, & \text{if } \lambda_0 - \tau > \delta_0. \end{cases} \tag{S29}$$

We define the extinction probability for purely bactericidal drugs ($\delta_T = \tau$ and $\lambda_T = 0$) as:

$$p_{E,\text{cidal}} = \begin{cases} 1, & \text{if } \lambda_0 \leq \delta_0 + \tau, \\ \frac{\delta_0+\tau}{\lambda_0}, & \text{if } \lambda_0 > \delta_0 + \tau. \end{cases} \tag{S30}$$

In Figure S37, we plot the colony formation probability for a single bacterium plated on agar ($p_C = 1 - p_E$) as a function of the treatment effect $\tau$, showing purely bacteriostatic (red) and purely bactericidal (blue) drugs.

#### S3 Filamentation Model

To model bacterial filamentation we discretize the cell volumes into $K$ classes indexed by $i \in \{1,2,\ldots,K\}$. Each class contains the bacterial density $b_i(t)$ of cells with volume

$$v_i = i\,\epsilon, \tag{S31}$$

where $\epsilon$ is a unit volume increment. We define the following rules for growth, division, and death of cells

- cells in class $i = 1$ cannot divide
- cells in class $i = K$ cannot grow
- cells in class $i = 1,\ldots,K-1$ shift from class $i$ to $i+1$ at rate $\gamma$
- cells in class $i = 2,\ldots,K$ divide at rate $\lambda$, resulting in a redistribution cells from class $i$ into smaller classes (e.g. if $i$ is even, two cells appear in class $i/2$; if $i$ is odd, one cell each appears in classes $\frac{i-1}{2}$ and $\frac{i+1}{2}$).
- cells in all classes die at rate $\delta$.

We collect the populations into a vector

$$\vec{b}(t) = \big(b_1(t), b_2(t), \ldots, b_K(t)\big)^{\mathsf{T}},$$

and write the dynamics as

$$\frac{d\vec{b}}{dt} = \lambda\Lambda\vec{b} + \gamma\Gamma\vec{b} - \delta\,\vec{b}, \tag{S32}$$

where $\Lambda$ and $\Gamma$ are transition matrices for division and growth, respectively. An example form for $\Gamma$ (volume acquisition) is

$$\Gamma = \begin{pmatrix} -1 & 0 & 0 & 0 & \cdots & 0 & 0 \\ 1 & -1 & 0 & 0 & \cdots & 0 & 0 \\ 0 & 1 & -1 & 0 & \cdots & 0 & 0 \\ \vdots & \vdots & \vdots & \vdots & \ddots & \vdots & \vdots \\ 0 & 0 & 0 & 0 & \cdots & 1 & 0 \end{pmatrix},$$

which shifts cells from class $i$ to $i+1$. An example $\Lambda$ (division) might be

$$\Lambda = \begin{pmatrix} 0 & 2 & 1 & 0 & \cdots & 0 \\ 0 & -1 & 1 & 2 & \cdots & 0 \\ 0 & 0 & -1 & 1 & \cdots & 0 \\ \vdots & \vdots & \vdots & \vdots & \ddots & \vdots \\ 0 & 0 & 0 & 0 & \cdots & -1 \end{pmatrix}.$$

We assume that the volume of the two new cells after division is identical to the original volume of the parent cell before division:

$$\sum_{i=1}^{K} i\,(\Lambda\vec{b})_i = 0. \tag{S33}$$

Furthermore the volume acquisition does not impact the number of cells:

$$\sum_{i=1}^{K} (\Gamma\vec{b})_i = 0. \tag{S34}$$

#### S3 a Population-level quantities.

We define the total bacterial density across all volume classes as

$$b(t) = \sum_{i=1}^{K} b_i(t), \tag{S35}$$

and *total biovolume density*:

$$V(t) = \epsilon \sum_{i=1}^{K} i\, b_i(t) = \sum_{i=1}^{K} v_i\, b_i(t). \tag{S36}$$

Then the *mean cell volume* is

$$v(t) = \frac{V(t)}{b(t)}. \tag{S37}$$

In the finite model, boundary effects arise because the smallest cells cannot divide and the largest cannot grow. In the continuum limit $\epsilon \to 0,\ K \to \infty$, these effects vanish and all cells experience uniform rates, so the following equalities hold:

$$\frac{db}{dt} = (\lambda - \delta)\, b, \tag{S38}$$

$$\frac{dV}{dt} = \epsilon\gamma\, b - \delta\, V, \tag{S39}$$

$$\frac{dv}{dt} = \epsilon\gamma - \lambda\, v. \tag{S40}$$

Setting $\frac{dv}{dt} = 0$ yields the equilibrium mean volume

$$v_{\mathrm{eq}} = \frac{\epsilon\gamma}{\lambda}. \tag{S41}$$

Integrating Equation S38 gives

$$b(t) = b_0 \exp\big((\lambda - \delta)t\big). \tag{S42}$$

Substituting this result into Equation S39 and integrating with the integrating factor $\exp(\delta t)$ yields the analytic solution for the total biovolume density accross all size classes:

$$V(t) = v_{\mathrm{eq}}\, b(t) + \big(V_0 - v_{\mathrm{eq}}\, b_0\big) \exp(-\delta t). \tag{S43}$$

Finally, writing $v_0 = V_0/b_0$, we obtain the analytical solution for the mean cell volume:

$$v(t) = v_{\mathrm{eq}} + \big(v_0 - v_{\mathrm{eq}}\big) \exp(-\lambda t). \tag{S44}$$

#### S3 b Parameter sensitivity.

We evaluated the impact of changes in the division rate, $\Delta\lambda \in [-1.5, 0.5]$, and the death rate, $\delta \in \{0, 2, 4\}$, on the difference between the luminescence-based rate and the true net growth rate. To this end, we set the treatment-free division rate to $\lambda_0 = 1.5$, the treatment-free death rate to $\delta_0 = 0$, the biovolume acquisition rate in the presence and absence of treatment to $\gamma = 150h^{-1}$ , the size of a volume increment to $\epsilon = 0.04\mu m^3$ , and volume-specific luminosity to $\theta = 0.015$. We simulated each parameter set for four hours using $K = 1000$. Furthermore we added the estimate based on luminescence if the first two hours of data are excluded.

#### S3 c Volume correction and parameter estimation

To correct the light signal for dynamic changes in biovolume, we combine two sources of information: (i) two morphology snapshots before ($v_{\mathrm{obs},0}$) and after 2 hours of treatment ($v_{\mathrm{obs,2h}}$), and (ii) the luminescence time series $I_{\mathrm{obs}}(t_i)$.
We first rewrite Equation S44 using the equilibrium–to–initial volume ratio $\alpha = v_{\mathrm{eq}}/v_0$ as:

$$v(t) = \alpha v_0 + (1 - \alpha)\, v_0\, \mathrm{e}^{-\lambda t}. \tag{S45}$$

To avoid fitting $\alpha$ as a free parameter, we insert $v_{\mathrm{obs},0}$ and $v_{\mathrm{obs,2h}}$ into Equation S45 to express $\alpha$ as a function of $\lambda$:

$$\alpha(\lambda) = \frac{v_{\mathrm{obs,2h}} - v_{\mathrm{obs},0}\mathrm{e}^{-2\lambda}}{v_{\mathrm{obs},0}(1 - \mathrm{e}^{-2\lambda})}, \qquad \alpha > 0. \tag{S46}$$

Assuming constant volume-specific luminosity, we insert this constrained $\alpha$ into Equation S11 to obtain:

$$I(t) = I_0\,\mathrm{e}^{(\lambda-\delta)t}\left[\alpha + (1-\alpha)\,\mathrm{e}^{-\lambda t}\right], \tag{S47}$$

with scale parameter $I_0 = \kappa\theta v_{\mathrm{obs},0} b_0 V_w$.
To estimate the division rate $\lambda$ and death rate $\delta$, we eliminate the nuisance parameter $I_0$ by defining

$$F_i := \mathrm{e}^{(\lambda-\delta)t_i}\left[\alpha + (1-\alpha)\,\mathrm{e}^{-\lambda t_i}\right], \tag{S48}$$

so that

$$\ln I(t_i) = \ln I_0 + \ln F_i. \tag{S49}$$

Minimizing the residual sum of squares over $\ln I_0$ yields the optimal

$$\ln I_0^* = \overline{\ln I_{\mathrm{obs}}} - \overline{\ln F}, \tag{S50}$$

where the overline denotes the sample mean across all $i$.
Minimizing the residual $\ln I(t_i) - \ln I_{\mathrm{obs}}(t_i)$ by substituting Eq. (S49) and Eq. (S50) yields the final loss function:

$$\mathcal{L}(\lambda,\delta) = \sum_{i=1}^{n}\left[\left(\ln F_i - \overline{\ln F}\right) - \left(\ln I_{\mathrm{obs}}(t_i) - \overline{\ln I_{\mathrm{obs}}}\right)\right]^2. \tag{S51}$$

To balance the dataset of observed specific volumes $v$, we sample 200 values per replicate and pool them. We then use the same bootstrapped light intensity datasets as in the main method for estimating $\psi_I$. For each bootstrap sample, we randomly pair one $v_{\mathrm{obs},0}$ and one $v_{\mathrm{obs},2\mathrm{h}}$ with one luminescence trajectory and minimize Eq. (S51) over the biologically plausible region:

$$0.01 \le \lambda \le 1.75, \qquad \delta \ge 0.$$

The remaining quantities $\psi = \lambda - \delta$, $\alpha(\lambda)$, and $I_0^*$ are computed algebraically.

### S4 Experiments

#### S4 a SOS experiment.

We conducted this experiment to test whether activating the SOS response by UV light would increase the specific luminosity and thereby explain the shallower decline of light intensity compared to the decline in CFU counts. This hypothesis rests on the possibility that the phage promoter driving the lux cassette up-regulates when the cell experiences stress.

To test this, we diluted three replicate overnight cultures 1:100 and grew them for approximately $1.5\,\mathrm{h}$ to mid-exponential phase. Each of the three exponential-phase cultures was split into two aliquots: one was assigned to UV treatment and placed in the upper half of a white 96-well plate (rows B–D), while the other served as an untreated control in the lower half (rows E–G) (Greiner, 655098).

We started the experiment by measuring luminescence and OD in the plate reader. Then we alternated between exposing the strains for repeated intervals to UV light in a cross-linker (Hoefer, UVC 500 crosslinker, at $10\,\frac{\mu\mathrm{J}}{\mathrm{cm}^2}$) in a temperature-regulated environment ($36.5\,°\mathrm{C}$); followed by luminescence and OD measurements. During UV exposure, we shielded the control samples by covering the lower half of the plate with a metal lid. The durations of UV treatment were 30 s, 1 min, 2 min, 4 min and 8 min.

OD increased in both UV-treated and control cultures; however, UV exposure visibly impaired OD growth compared to the controls (Figure S21a). We normalized the luminescence signals (Figure S21b) by dividing through the OD signal, resulting in the OD-normalized light intensity $\omega$ (Figure S21c). We observed that the OD-normalized light intensity of UV-treated cultures falls with the duration of treatment compared to the OD-normalized light intensity of the controls ($\omega_{\mathrm{UV}} - \omega_{\mathrm{ctrl}}$; see Figure S21d; t-test, $p = 3 \cdot 10^{-5}$ for the last timepoint).

Based on these results, we find it unlikely that upregulation of the promoter explains the shallower decline of light intensity compared to that of CFU count. However, we cannot exclude the possibility that this result does not hold if the SOS response is triggered by another mechanism.

#### S4 b Morphology evaluation.

We analyzed the microscopy images in several steps. First, we manually applied lower and upper thresholds to the red and green channel to enhance the visual contrast (Figure S22–Figure S34). Next, we used "Ilastik-1.4.0" to infer the probability that each pixel in the thresholded green channel image belonged to a bacterium. Ilastik employs a neural network trained directly on the microscopy images.

Subsequently, we used a Python script ([5]) to convert these probabilities into markers representing individual bacteria. Misidentified markers were manually excluded, e.g., if they only partially covered a bacterium or covered multiple overlapping bacteria. For each marker, we fitted a spline through its center, providing the spline length $l_s$. We then optimized the radius $r_s$ by maximizing the marker area within a distance $r_s$ from the spline while minimizing the area within $r_s$ that did not belong to the marker.

Using these parameters, we calculated for each bacterium the length $l_b = l_s + 2 \cdot r_s$, width $w_b = 2 \cdot r_s$, and volume $v = 2 \cdot \pi \cdot r_s^2 \cdot l_s + \frac{4}{3} \cdot \pi \cdot r_s^3$.To create a volume distribution for each treatment, we resampled the fitted volume estimates from each image (replicate) 200 times with replacement, preserving the original sample size, and aggregated the resulting datasets. Based on these distributions, we determined whether cells were significantly filamented using the significance criterion described in the methods section of the main paper.

#### S4 c Antimicrobial peptide deactivation experiment.

To assess whether pexiganan-treated bacteria continued to die in a 1:100 diluted PBS environment, we exposed exponential-phase cultures to $16\,\frac{\mu\mathrm{g}}{\mathrm{mL}}$ pexiganan for $1\,\mathrm{min}$. Following this treatment, $10\,\mu\mathrm{L}$ of each culture was diluted in $990\,\mu\mathrm{L}$ of PBS supplemented with 0, 1, 10, or $100\,\mathrm{mM}$ $CaCl_2$ or $MgCl_2$. Every $45\,\mathrm{min}$, we sampled from each diluted culture and plated $10\,\mu\mathrm{L}$ aliquots using the automated plating method described in the methods section of the main paper.

We observed a substantial effect of both supplements on the measured bacterial density (Figure S38a, b). Increasing the supplement concentration consistently resulted in higher bacterial densities, indicating that the supplemented ions reduce bacterial killing. Most data points for the unsupplemented medium resulted in empty agar plates. The highest CFU count was observed for strains diluted in $100\,\mathrm{mM}$ $MgCl_2$ (Table S4).

#### S4 d Manual pexiganan time-kill curve experiment.

In this setup, we captured four time points within $5\,\mathrm{min}$ using CFU plus the pre-treatment bacterial density. The experiment was performed manually using the traditional CFU plating method. Round agar plates (Sarstedt, 82.1473.001) containing $25\,\mathrm{mL}$ of agar were used, and $100\,\mu\mathrm{L}$ of each dilution was plated. This approach increases sensitivity by using $100\,\mu\mathrm{L}$, rather than $10\,\mu\mathrm{L}$, for plating.

Cultures were treated in a 96-deepwell plate (Greiner, 780285) by adding $100\,\mu\mathrm{L}$ of a 10x stock to $900\,\mu\mathrm{L}$ of exponential-phase culture. Samples were taken directly from the deepwell plate, and two simultaneous dilution series were prepared in a 96-well

plate (Greiner, 655101) at each time point to halt the killing. One dilution series was prepared in pure PBS, and the other in PBS supplemented with $100\,\mathrm{mM}$ $MgCl_2$.
For this experiment, we plated three dilutions (factors of 100, 1,000, and 10,000) and counted the colonies on all plates. We observed that when PBS without $MgCl_2$ was used, higher dilution factors led to higher CFU estimates (Figure S39). This discrepancy diminished over time, in parallel with a weakening of the observed kill rate. In contrast, when the dilution medium was supplemented with $MgCl_2$, we did not observe this effect.

#### S4 e Supernatant experiment.

During the measured pexiganan kill curve described above, we observed a steep decline in CFU counts, followed by a nearly constant plateau. Two non-exclusive explanations may account for this observed decrease in killing: (i) the surviving bacteria are persisters or resistant to the AMP, or (ii) the AMP molecules become deactivated, leaving the supernatant without killing activity.
To investigate the supernatant's remaining bactericidal effects, we conducted a multi-step experiment:
**Step 1: preparation.** Three overnight (O/N) cultures were diluted 1:100 in LB and incubated at $37\,^{\circ}\mathrm{C}$ with shaking for $2\,\mathrm{h}$. From each culture, we took three samples: one to measure the bacterial density before treatment, the second (1 ml) to accumulate pure bacteria for supernatant exposure, and the third $1.35\,\mathrm{mL}$ was reserved for supernatant production and measuring the initial kill rate.
**Step 2: purification of bacteria.** To purify bacteria we pelleted the previously collected $1\,\mathrm{mL}$ bacterial aliquots at $3000\,\times\mathrm{g}$ for $5\,\mathrm{min}$, discarded the supernatant and stored them in the fridge.
**Step 3: initial killing and supernatant production.** To generate the supernatant, each $1.35\,\mathrm{mL}$ sample was treated with a $160\,\frac{\mathrm{\mu g}}{\mathrm{mL}}$ pexiganan stock at a 1:10 ratio, yielding a final concentration of $16\,\frac{\mathrm{\mu g}}{\mathrm{mL}}$. After $5\,\mathrm{min}$, we plated samples for CFU counts and centrifuged the remaining culture at maximum speed for $2\,\mathrm{min}$ to remove cellular debris. Plate counts of these treated samples revealed rapid bacterial killing (rate of CFU count change $\psi_{\mathrm{CFU}} \approx -46$ per hour) (Figure S40, Table S5). We collected $1\,\mathrm{mL}$ of the clarified supernatant for the subsequent exposure experiment.
**Step 4: supernatant killing.** In that final step we dissolved the pelleted bacteria (from step 2) in the supernatant collected during step 3. After another $5\,\mathrm{min}$ incubation, we plated samples (dilutions 1:100, 1:1,000, and 1:10,000) to estimate bacterial density. No significant killing was observed, indicating that the supernatant alone no longer exhibited bactericidal activity—supporting explanation (ii), without rejecting (i).
Our current hypothesis is that AMP molecules bind to the surface of intact bacteria and to newly exposed targets from lysed bacteria, thereby becoming deactivated. Thus, cells that survive the initial kill phase may have an increased chance of continued survival. Possible explanations for why specific bacteria survive this phase include reduced surface area due to clumping or adhesion to well walls, smaller cell size, and other factors that might confer protection, potentially related to the cell cycle.

#### S5 Tables

**Table S1:** Drugs used in this study, their MICs, working concentrations, and stock solvents. In the MIC column, we report the highest concentration of the dilution series (numerator) and the maximum inhibiting dilution (denominator). Kanamycin ($50\ \frac{\mu g}{mL}$) was used as the selection marker for the *lux* operon.

| drug | MIC [$\frac{\mu g}{mL}$] | cwork [$\frac{\mu g}{mL}$] | cwork [MIC] | solvent | supplier |
|---|---|---|---|---|---|
| Amoxicillin | $\frac{10}{4} = 2.50$ | 25 | 10 | DMSO | Sigma, A8523 |
| Ampicillin | $\frac{100}{128} = 0.78$ | 10, 2 | 12.8, 2.56 | water | Sigma, A9518 |
| Cefepime | $\frac{4}{256} = 0.02$ | 0.15 | 9.6 | DMSO | ThermoFisher, J66237 |
| Ceftazidime | $\frac{4}{64} = 0.06$ | 0.62 | 10 | DMSO | Sigma, PHR1847 |
| Cefuroxime | $\frac{4}{2} = 2.00$ | 16 | 8 | water | Sigma, C4417 |
| Chloramphenicol | $\frac{128}{64} = 2.00$ | 20 | 10 | DMSO | Sigma, C0378 |
| Ciprofloxacin | $\frac{1}{128} = 0.01$ | 0.08 | 10 | water | Sigma, 17850 |
| Colistin | $\frac{50}{64} = 0.78$ | 1.6 | 2.05 | water | Sigma, C4461 |
| Doripenem | $\frac{4}{256} = 0.02$ | 0.16 | 10.24 | water | VWR, ACRO463870010 |
| Fosfomycin | $\frac{4}{4} = 1.00$ | 8 | 8 | water | VWR, APOSBIM0107 |
| Imipenem | $\frac{5}{64} = 0.08$ | 1 | 12.8 | water | Sigma, PHR1796 |
| Mecilinam | $\frac{10}{128} = 0.08$ | 0.78 | 10 | DMSO | Sigma, 33447 |
| Meropenem | $\frac{5}{512} = 0.01$ | 0.1 | 10.24 | water | Sigma, PHR1772 |
| Penicillin | $\frac{500}{32} = 15.62$ | 100 | 6.4 | water | Roth, HP48.2 |
| Pexiganan | $\frac{64}{32} = 2.00$ | 8, 16 | 4, 8 | water | Sigma, SML3787 |
| Piperacillin | $\frac{10}{16} = 0.62$ | 6.25 | 10 | DMSO | Sigma, J66419 |
| Polymyxin B | $\frac{50}{32} = 1.56$ | 2.5 | 1.6 | water | Roth, 0235.1 |
| Rifampicin | $\frac{40}{16} = 2.50$ | 25 | 10 | DMSO | Sigma, R3501 |
| Tetracycline | $\frac{10}{32} = 0.31$ | 3.12 | 10 | DMSO | Sigma, T3383 |
| Trimethoprim | $\frac{10}{128} = 0.08$ | 0.78 | 10 | DMSO | Sigma, T7883 |

**Table S2:** Point estimates and 95% percentile intervals of $\psi_{\mathrm{CFU}}$, $\psi_I$, $\psi_I^*$, and $\psi_J$ for different treatments. Sig$_X$ indicates whether the rate of change of signal $X \in \{I, I^*, J\}$ differs significantly (*) from the distribution of $\psi_{\mathrm{CFU}}$, or not (n.s.), based on the significance criterion defined in the Methods section. Estimates are based on data from the CFU-luminescence assays (see Methods).

| | $\psi_{\mathrm{CFU}}$ $[\frac{1}{\mathrm{h}}]$ | $\psi_I$ $[\frac{1}{\mathrm{h}}]$ | sig$_I$ | $\psi_I^*$ $[\frac{1}{\mathrm{h}}]$ | sig$_{I^*}$ | $\psi_J$ $[\frac{1}{\mathrm{h}}]$ | sig$_J$ |
|---|---|---|---|---|---|---|---|
| **ampicillin** 10 $\frac{\mu g}{mL}$ | -3.23 [-4.11, -2.54] | -2.41 [-2.91, -2.11] | * | | | -2.63 [-3.31, -2.29] | n.s. |
| **ampicillin** 2 $\frac{\mu g}{mL}$ | -0.81 [-1.19, -0.37] | -0.30 [-0.41, -0.22] | * | | | | |
| **amoxicillin** 25 $\frac{\mu g}{mL}$ | -2.15 [-2.42, -1.88] | -2.04 [-2.21, -1.86] | n.s. | | | | |
| **cefepime** 0.15 $\frac{\mu g}{mL}$ | -1.17 [-1.50, -0.85] | -0.47 [-0.74, -0.24] | * | -0.88 [-1.13, -0.65] | * | | |
| **ceftazidime** 0.62 $\frac{\mu g}{mL}$ | -0.88 [-1.06, -0.71] | -0.09 [-0.24, 0.04] | * | | | -0.38 [-0.52, -0.15] | * |
| **cefuroxime** 16 $\frac{\mu g}{mL}$ | -1.59 [-2.41, -0.99] | -1.41 [-1.64, -1.25] | n.s. | -1.62 [-1.92, -1.34] | n.s. | | |
| **chloramphenicol** 20 $\frac{\mu g}{mL}$ | 0.05 [-0.11, 0.20] | 0.04 [-0.03, 0.10] | n.s. | | | | |
| **ciprofloxacin** 0.078 $\frac{\mu g}{mL}$ | -1.96 [-2.29, -1.64] | 0.83 [0.44, 1.17] | * | | | 0.48 [0.09, 0.90] | * |
| **colistin** 1.6 $\frac{\mu g}{mL}$ | -1.00 [-1.45, -0.52] | -0.93 [-1.26, -0.66] | n.s. | | | | |
| **doripenem** 0.16 $\frac{\mu g}{mL}$ | -0.54 [-0.83, -0.25] | -0.24 [-0.37, -0.15] | * | -0.37 [-0.53, -0.26] | * | | |
| **fosfomycin** 8 $\frac{\mu g}{mL}$ | -1.22 [-1.65, -0.83] | -1.12 [-1.37, -0.93] | n.s. | | | | |
| **imipenem** 1 $\frac{\mu g}{mL}$ | -1.42 [-2.06, -0.72] | 0.00 [-0.16, 0.14] | * | -0.32 [-0.45, -0.22] | * | | |
| **mecillinam** 0.78 $\frac{\mu g}{mL}$ | -0.32 [-0.57, -0.10] | 0.02 [-0.17, 0.17] | * | -0.34 [-0.47, -0.20] | n.s. | | |
| **meropenem** 0.1 $\frac{\mu g}{mL}$ | -1.85 [-2.36, -1.32] | -0.16 [-0.47, 0.10] | * | | | -0.55 [-0.80, -0.21] | * |
| **penicillin** 100 $\frac{\mu g}{mL}$ | -2.17 [-2.78, -1.66] | -2.20 [-2.53, -1.94] | n.s. | | | | |
| **piperacillin** 6.25 $\frac{\mu g}{mL}$ | -0.26 [-0.51, 0.02] | 0.09 [-0.01, 0.19] | * | | | | |
| **polymyxinB** 2.5 $\frac{\mu g}{mL}$ | -1.45 [-1.73, -1.22] | -1.41 [-1.87, -1.07] | n.s. | | | | |
| **rifampicin** 25 $\frac{\mu g}{mL}$ | -0.23 [-0.40, 0.04] | -0.15 [-0.23, -0.07] | n.s. | | | | |
| **tetracycline** 3.125 $\frac{\mu g}{mL}$ | -0.06 [-0.12, 0.00] | -0.01 [-0.07, 0.05] | n.s. | | | | |
| **trimethoprim** 0.78 $\frac{\mu g}{mL}$ | -0.61 [-0.76, -0.47] | 0.48 [0.38, 0.59] | * | | | 0.34 [0.18, 0.51] | * |
| **pexiganan** 8 $\frac{\mu g}{mL}$ | -44.59 [-66.19, -20.08] | -45.95 [-48.21, -43.34] | n.s. | | | | |
| **pexiganan** 16 $\frac{\mu g}{mL}$ | -61.63 [-93.40, 1.26] | -60.27 [-63.06, -56.42] | n.s. | | | | |

**Table S3:** Bootstrapped 95% confidence intervals and point estimates for the length, width, and volume of cells after 2 h of treatment, estimated from microscopy images. Significance was assessed by comparing the confidence intervals of cell volumes for each antibiotic treatment to the untreated control (`control_2h`), as described in the Methods section of the main paper.

| | L_lower | L_upper | L_mean | W_lower | W_upper | W_mean | V_lower | V_upper | V_mean | V_sig |
|---|---|---|---|---|---|---|---|---|---|---|
| control_2h | 2.28 | 4.92 | 3.41 | 0.77 | 1.4 | 1.09 | 2 | 6.51 | 3.89 | ref. |
| amoxicillin | 2.64 | 10.77 | 5.09 | 0.47 | 1.37 | 1.17 | 1.93 | 9.67 | 6.21 | n.s. |
| ampicillin | 3.87 | 27.14 | 12.36 | 0.93 | 1.67 | 1.27 | 5.7 | 30.39 | 16.67 | significant |
| ceftazidime | 32.05 | 68.69 | 54.54 | 0.67 | 1.22 | 0.95 | 11.52 | 80.86 | 40.42 | significant |
| ciprofloxacin | 10.88 | 36.82 | 21.77 | 0.7 | 1.27 | 1.03 | 6.35 | 45.7 | 19.55 | significant |
| colistin | 2.19 | 4.53 | 3.14 | 0.6 | 1.59 | 1.07 | 1 | 8.32 | 3.71 | n.s. |
| fosfomycin | 1.77 | 5.32 | 3.18 | 0.69 | 1.15 | 0.93 | 1.09 | 4.75 | 2.65 | n.s. |
| meropenem | 3.44 | 10.61 | 5.9 | 1.13 | 3.71 | 2.26 | 5.33 | 71.01 | 31.11 | significant |
| rifampicin | 2.41 | 8.38 | 4.77 | 0.59 | 1.4 | 0.98 | 1.15 | 9.44 | 4.32 | n.s. |
| tetracycline | 2.43 | 7.7 | 4.71 | 0.65 | 1.71 | 1.08 | 1.29 | 13.4 | 5.25 | n.s. |
| trimethoprim | 4.67 | 30.74 | 12.19 | 0.65 | 1.29 | 0.93 | 2.64 | 19.79 | 9.05 | significant |

**Table S4:** Estimated group means and $95\%$ confidence intervals from an ordinary least squares (OLS) model fitted to log-transformed CFU data, collected from the first sampled time point after diluting pexiganan-treated strains in supplemented PBS. Grouping is based on the supplement ($CaCl_2$ or $MgCl_2$) and concentration ($0\,\mathrm{mM}$ to $100\,\mathrm{mM}$). Confidence intervals were computed using heteroscedasticity-consistent standard errors (HC3). The compact letter display (cld) indicates groups that are not significantly different by sharing a common letter, based on mutual inclusion of their $95\%$ confidence intervals.

| group | mean | ci_lower | ci_upper | cld |
|---|---|---|---|---|
| $CaCl_2$ (0 mM) | -0.00 | -0.00 | 0.00 | a |
| $CaCl_2$ (1 mM) | 2.77 | -0.56 | 6.09 | b |
| $CaCl_2$ (10 mM) | 5.06 | 4.92 | 5.20 | c |
| $CaCl_2$ (100 mM) | 5.85 | 5.38 | 6.31 | d |
| $MgCl_2$ (0 mM) | -0.00 | -0.00 | 0.00 | a |
| $MgCl_2$ (1 mM) | 2.77 | -0.56 | 6.09 | b |
| $MgCl_2$ (10 mM) | 5.31 | 5.20 | 5.42 | e |
| $MgCl_2$ (100 mM) | 6.36 | 6.27 | 6.46 | f |

**Table S5:** Comparison of kill rates $[h^{-1}]$ between cultures treated with pexiganan ($16\,\frac{\mathrm{\mu g}}{\mathrm{mL}}$) for 5 minutes and cultures exposed to the supernatant collected after the kill assay. The two rates differ significantly; the confidence interval of the rate of change of CFU in the supernatant includes zero.

| | mean | lower | upper |
|---|---|---|---|
| experiment | | | |
| pexiganan | -46.980603 | -63.153906 | -35.797836 |
| supernatant | 1.281551 | -2.187859 | 5.111196 |

**Table S6:** Primer sequences used for $\lambda$-red mediated integration of the *luxCDABE* operon into *E. coli*. Lowercase letters indicate homology regions binding to the *lux* operon on the plasmid; uppercase letters indicate chromosomal homology regions at the integration site.

| Primer name | Sequence (5' $\rightarrow$ 3') |
|---|---|
| forward primer | CGGTACGGCTGACCATCGGGTGCCAGTGCGGGAGTTTCGT**acccagtaaggcagcggtatc** |
| reverse primer | AGTCAGCGATATCCATTTTCGCGAATCCGGAGTGTAAGAA**taggtctagggcggcgga** |

#### S6 Figures

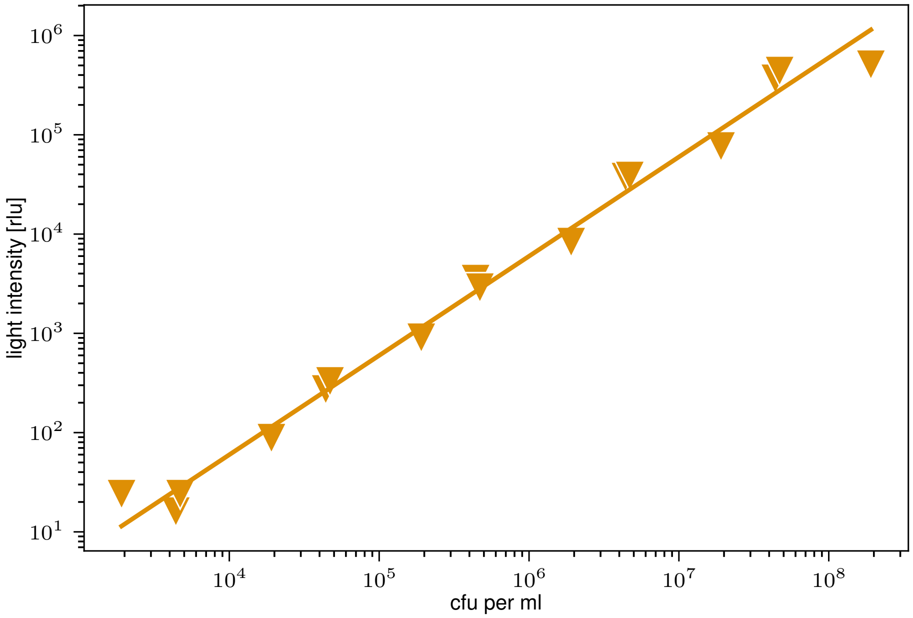


**Figure S1:** Light intensity scales linearly with bacterial density. Serial tenfold dilutions of bacterial cultures were prepared in a 384-well white microplate, and luminescence was measured immediately. Linear regression of the luminescence signal against bacterial density (CFU) yielded a conversion factor of $m_{\mathrm{fit}} = 0.006\,\mathrm{rlu} \cdot \mathrm{ml} \cdot \mathrm{CFU}^{-1}$. The high correlation ($R^2 = 0.987$ in log-log space) confirms a linear relationship between luminescence and bacterial density.

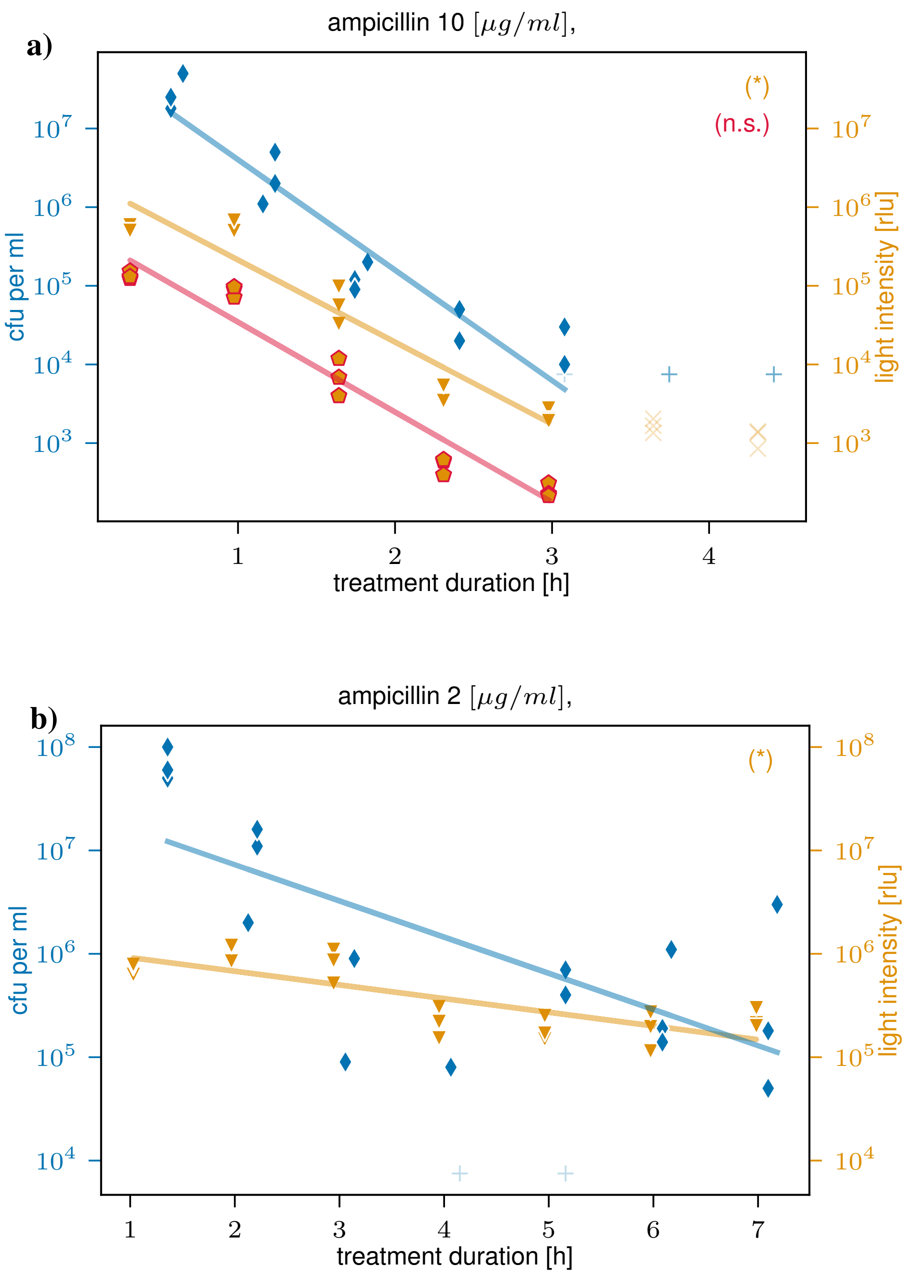


**Figure S2:** Ampicillin. The CFU signal is shown as blue diamonds and the light intensity as orange triangles for data points above their respective detection limits ($10^4\,\mathrm{CFU/mL}$ for CFU and 20 rlu for luminescence). Lines represent log-linear fits to the corresponding signals. The morphology-corrected luminescence signal is shown as orange pentagons with red frames, and the corresponding rate fit indicated by a red line. '+' indicates data points below detection limit or otherwise excluded (for CFU, symbolically plotted at $10^4\,\mathrm{CFU/mL}$ to visualize missing data), and If a signal ends early (due to dropping below its detection limit), the corresponding data point of the other signal was cut to the same endpoint for a consitent comparison. These excluded data points are marked as 'X'.

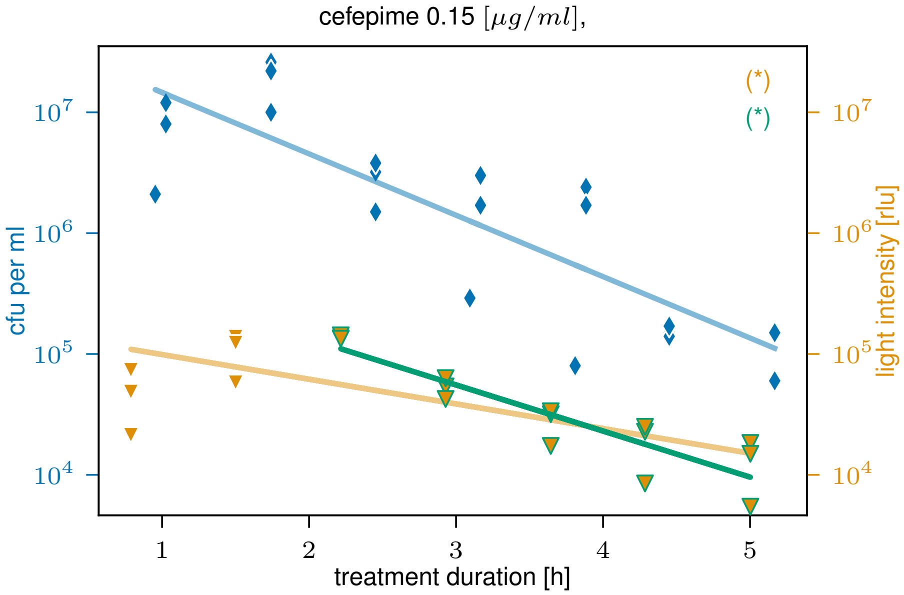


**Figure S3:** Cefepime. The CFU signal is shown as blue diamonds and the light intensity as orange triangles for data points above their respective detection limits ($10^4$ CFU/mL for CFU and 20 rlu for luminescence). Lines represent log-linear fits to the corresponding signals. Green-framed data points and corresponding green fit lines indicate analyses excluding early data points until the first peak.

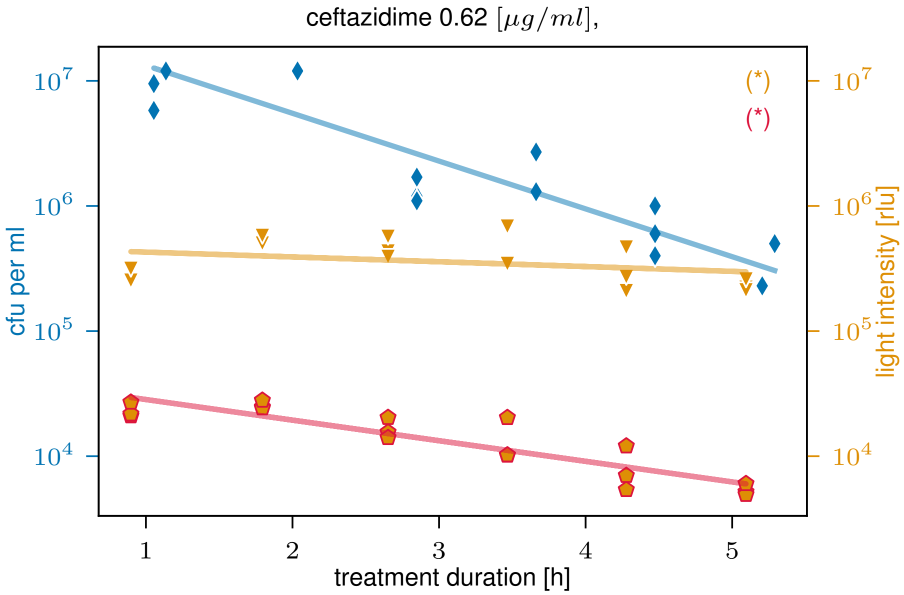


**Figure S4:** Ceftazidime. The CFU signal is shown as blue diamonds and the light intensity as orange triangles for data points above their respective detection limits ($10^4$ CFU/mL for CFU and 20 rlu for luminescence). Lines represent log-linear fits to the corresponding signals. The morphology-corrected luminescence signal is shown as orange pentagons with red frames, and the corresponding rate fit indicated by a red line.

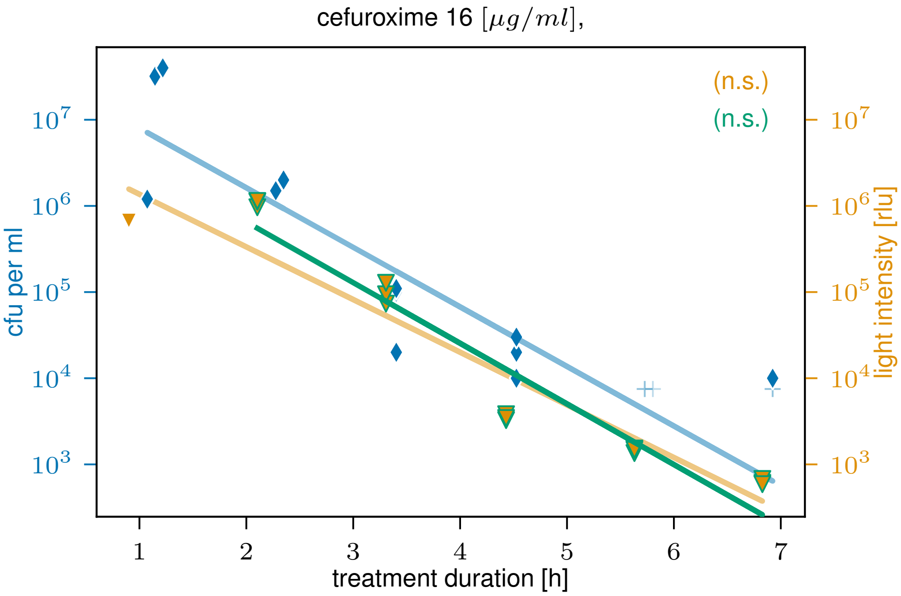


**Figure S5:** Cefuroxime. The CFU signal is shown as blue diamonds and the light intensity as orange triangles for data points above their respective detection limits ($10^4$ CFU/mL for CFU and 20 rlu for luminescence). Lines represent log-linear fits to the corresponding signals. '+' indicates data points below detection limit or otherwise excluded (for CFU, symbolically plotted at $10^4$ CFU/mL to visualize missing data).

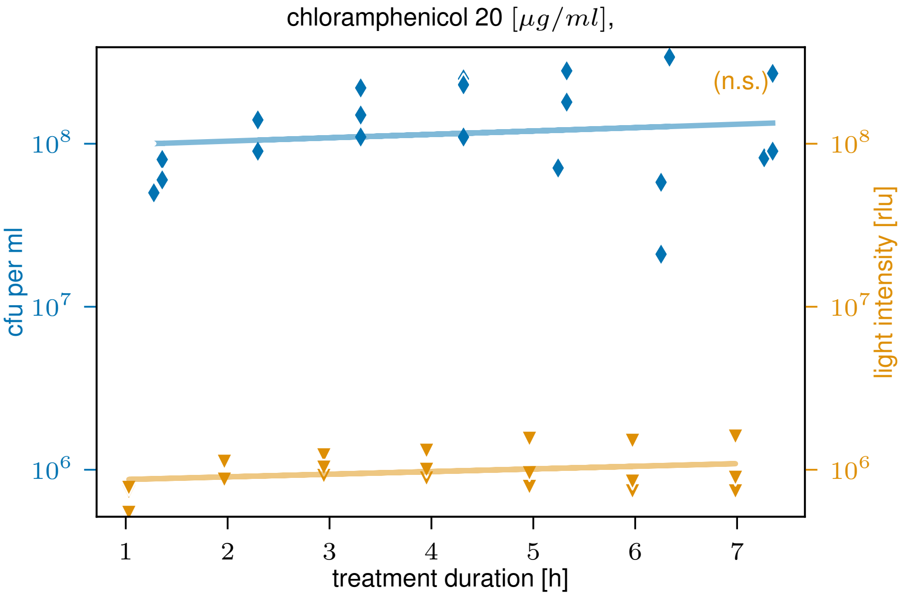


**Figure S6:** Chloramphenicol. The CFU signal is shown as blue diamonds and the light intensity as orange triangles for data points above their respective detection limits ($10^4$ CFU/mL for CFU and 20 rlu for luminescence). Lines represent log-linear fits to the corresponding signals.

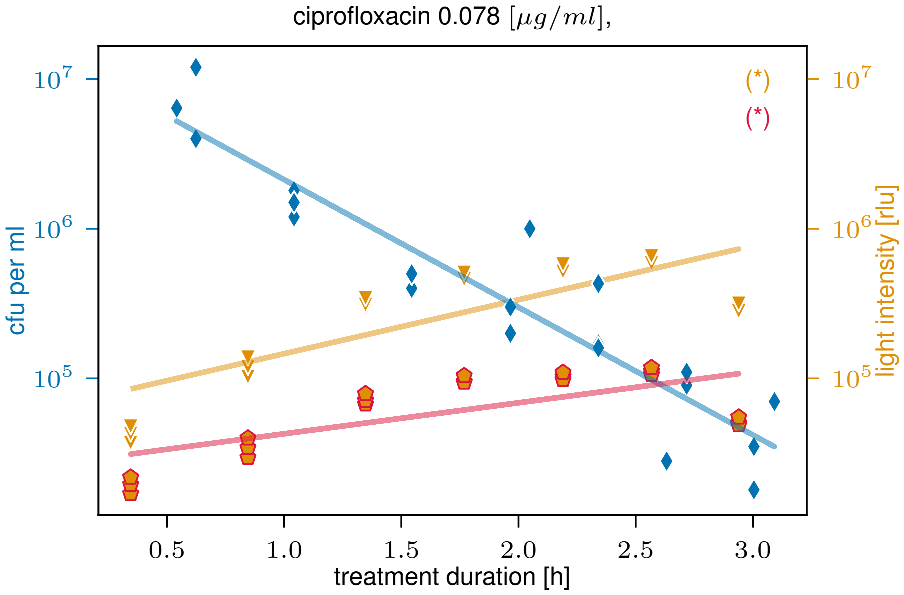


**Figure S7:** Ciprofloxacin. The CFU signal is shown as blue diamonds and the light intensity as orange triangles for data points above their respective detection limits ($10^4$ CFU/mL for CFU and 20 rlu for luminescence). Lines represent log-linear fits to the corresponding signals. The morphology-corrected luminescence signal is shown as orange pentagons with red frames, and the corresponding rate fit indicated by a red line.

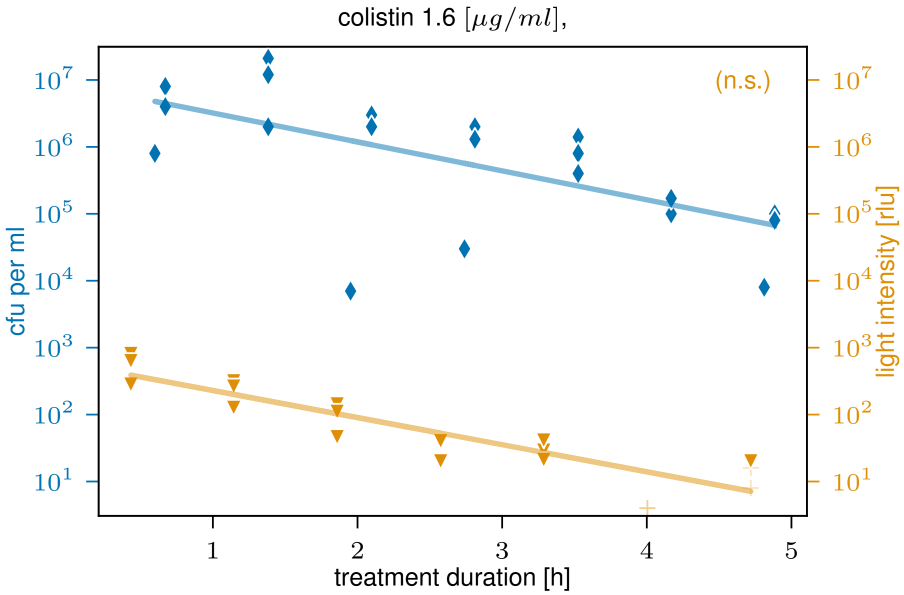


**Figure S8:** Colistin. The CFU signal is shown as blue diamonds and the light intensity as orange triangles for data points above their respective detection limits ($10^4$ CFU/mL for CFU and 20 rlu for luminescence). Lines represent log-linear fits to the corresponding signals. '+' indicates data points below detection limit or otherwise excluded (for CFU, symbolically plotted at $10^4$ CFU/mL to visualize missing data)

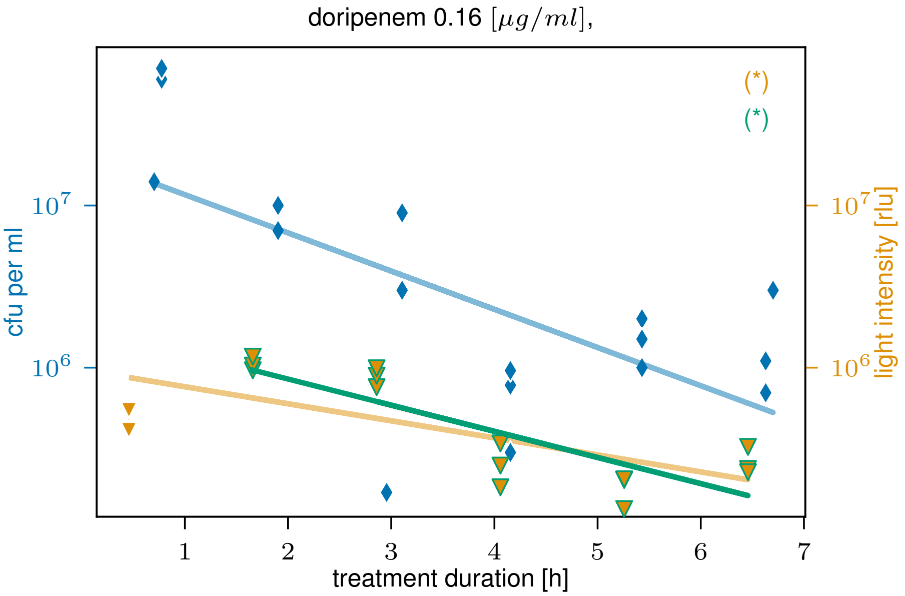


**Figure S9:** Doripenem. The CFU signal is shown as blue diamonds and the light intensity as orange triangles for data points above their respective detection limits ($10^4$ CFU/mL for CFU and 20 rlu for luminescence). Lines represent log-linear fits to the corresponding signals. Green-framed data points and corresponding green fit lines indicate analyses excluding early data points until the first peak.

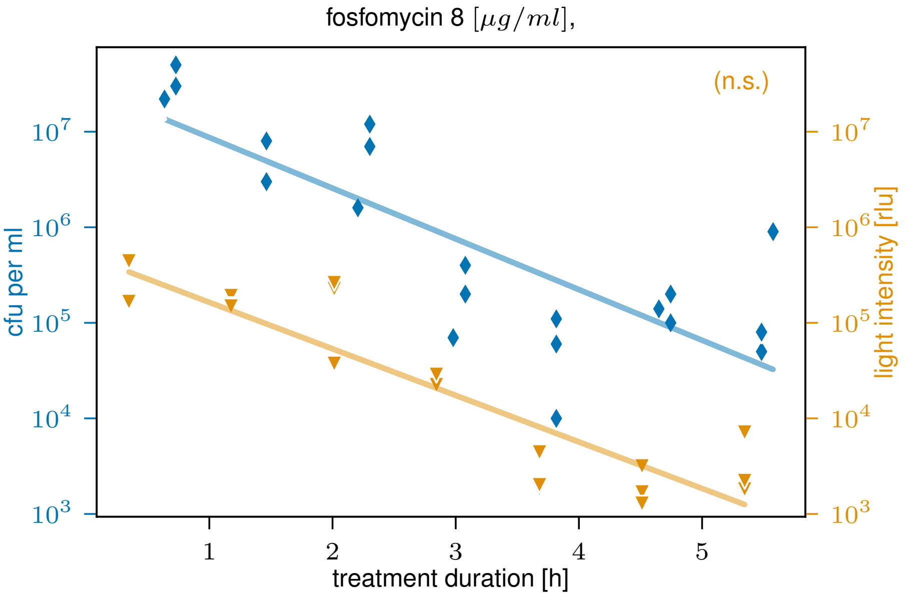


**Figure S10:** Fosfomycin. The CFU signal is shown as blue diamonds and the light intensity as orange triangles for data points above their respective detection limits ($10^4$ CFU/mL for CFU and 20 rlu for luminescence). Lines represent log-linear fits to the corresponding signals.

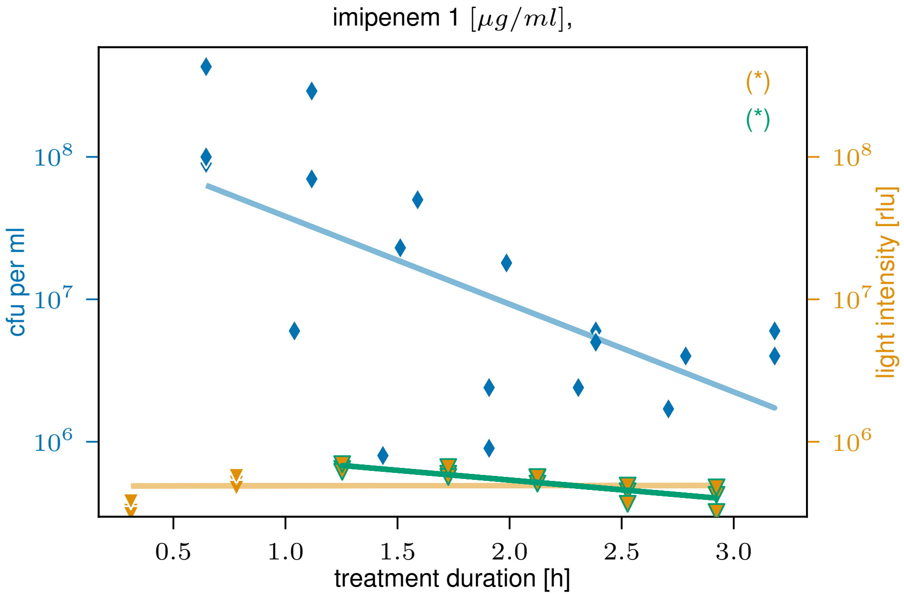


**Figure S11:** Imipenem. The CFU signal is shown as blue diamonds and the light intensity as orange triangles for data points above their respective detection limits ($10^4$ CFU/mL for CFU and 20 rlu for luminescence). Lines represent log-linear fits to the corresponding signals. '+' indicates data points below detection limit or otherwise excluded (for CFU, symbolically plotted at $10^4$ CFU/mL to visualize missing data).

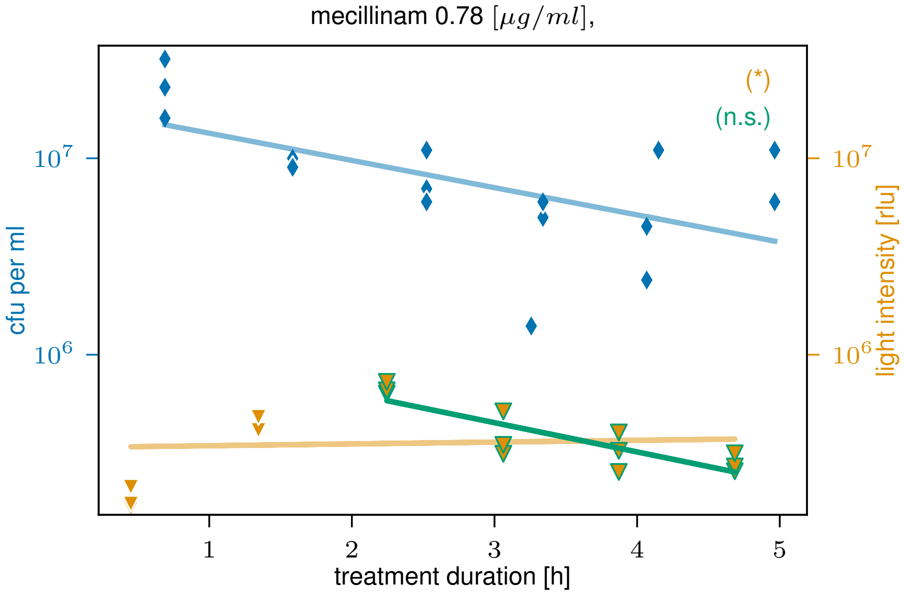


**Figure S12:** Mecillinam. The CFU signal is shown as blue diamonds and the light intensity as orange triangles for data points above their respective detection limits ($10^4$ CFU/mL for CFU and 20 rlu for luminescence). Lines represent log-linear fits to the corresponding signals.

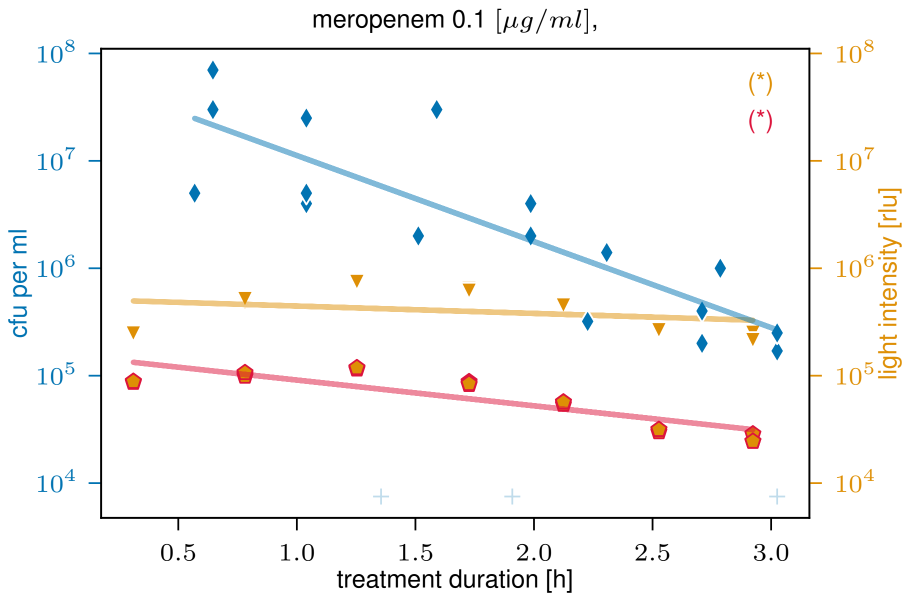


**Figure S13:** Meropenem. The CFU signal is shown as blue diamonds and the light intensity as orange triangles for data points above their respective detection limits ($10^4$ CFU/mL for CFU and 20 rlu for luminescence). Lines represent log-linear fits to the corresponding signals. The morphology-corrected luminescence signal is shown as orange pentagons with red frames, and the corresponding rate fit indicated by a red line. '+' indicates data points below detection limit or otherwise excluded (for CFU, symbolically plotted at $10^4$ CFU/mL to visualize missing data).

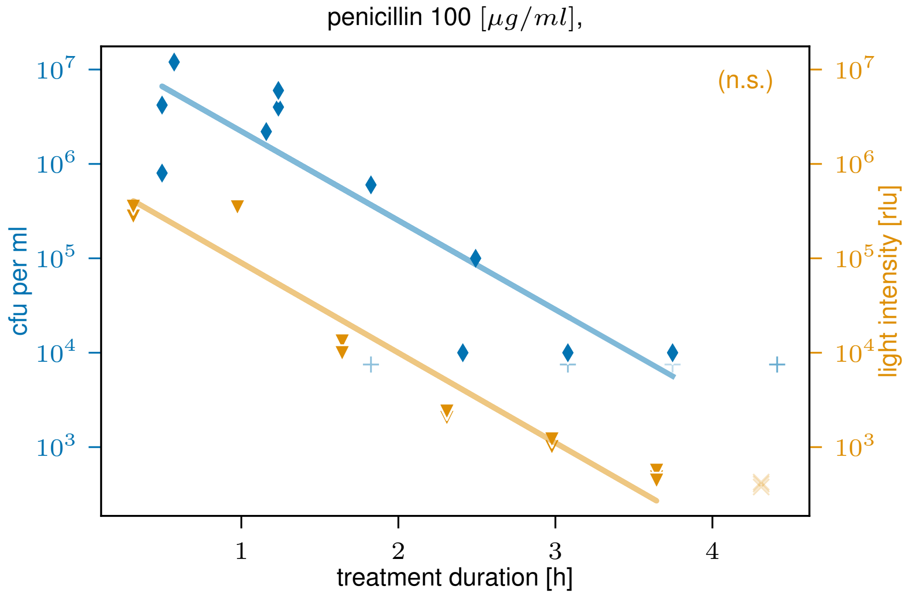


**Figure S14:** Penicillin. The CFU signal is shown as blue diamonds and the light intensity as orange triangles for data points above their respective detection limits ($10^4$ CFU/mL for CFU and 20 rlu for luminescence). Lines represent log-linear fits to the corresponding signals. '+' indicates data points below detection limit or otherwise excluded (for CFU, symbolically plotted at $10^4$ CFU/mL to visualize missing data), and If a signal ends early (due to dropping below its detection limit), the corresponding data point of the other signal was cut to the same endpoint for a consitent comparison. These excluded data points are marked as 'X'.

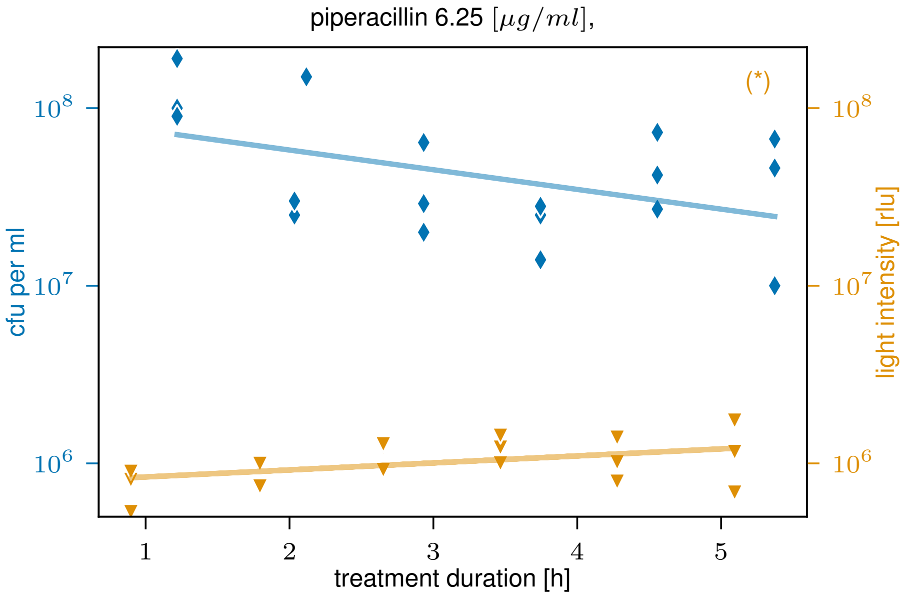


**Figure S15:** Piperacillin. The CFU signal is shown as blue diamonds and the light intensity as orange triangles for data points above their respective detection limits ($10^4$ CFU/mL for CFU and 20 rlu for luminescence). Lines represent log-linear fits to the corresponding signals.

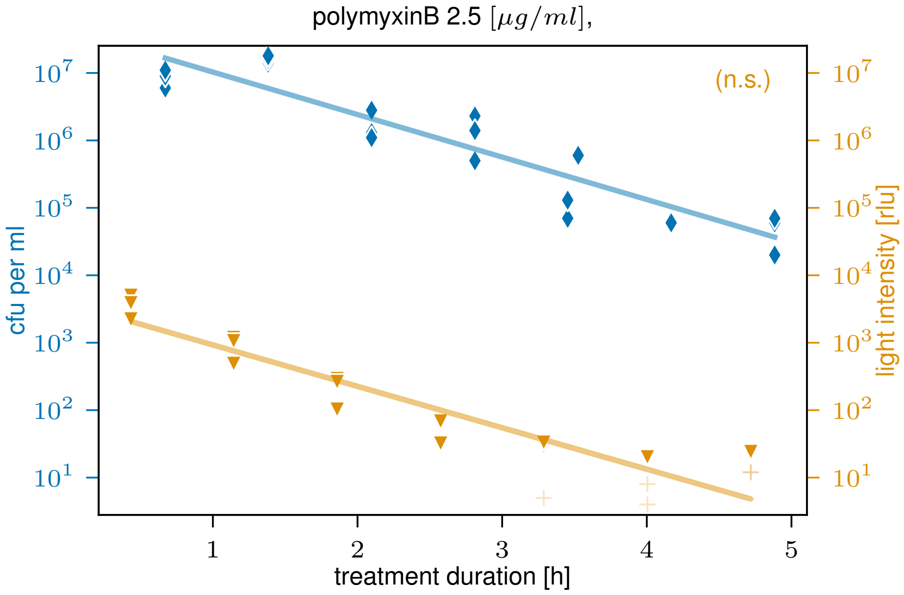


**Figure S16:** Polymyxin B. The CFU signal is shown as blue diamonds and the light intensity as orange triangles for data points above their respective detection limits ($10^4$ CFU/mL for CFU and 20 rlu for luminescence). Lines represent log-linear fits to the corresponding signals. '+' indicates data points below detection limit or otherwise excluded (for CFU, symbolically plotted at $10^4$ CFU/mL to visualize missing data).

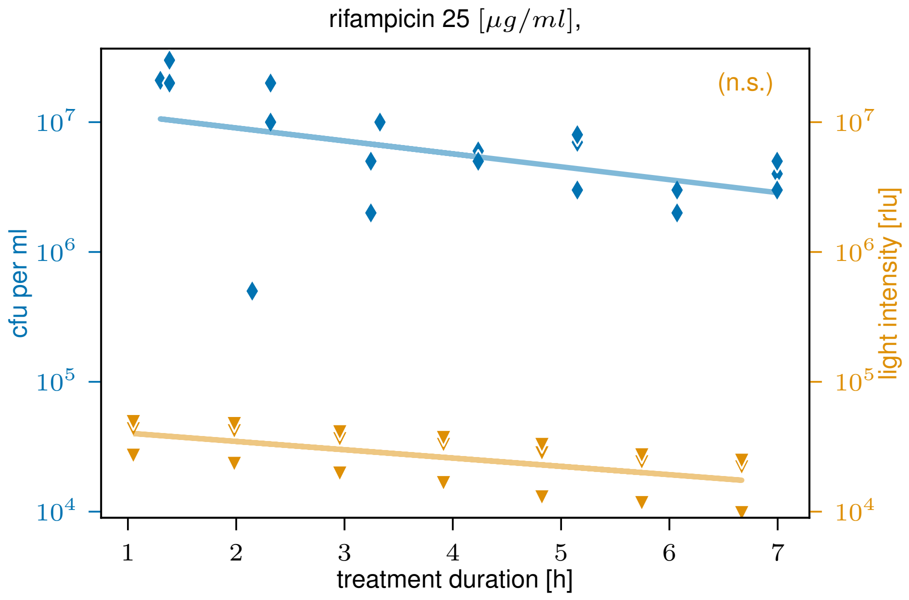


**Figure S17:** Rifampicin ($25\,\mu g/mL$). The CFU signal is shown as blue diamonds and the light intensity as orange triangles for data points above their respective detection limits ($10^4$ CFU/mL for CFU and 20 rlu for luminescence). Lines represent log-linear fits to the corresponding signals.

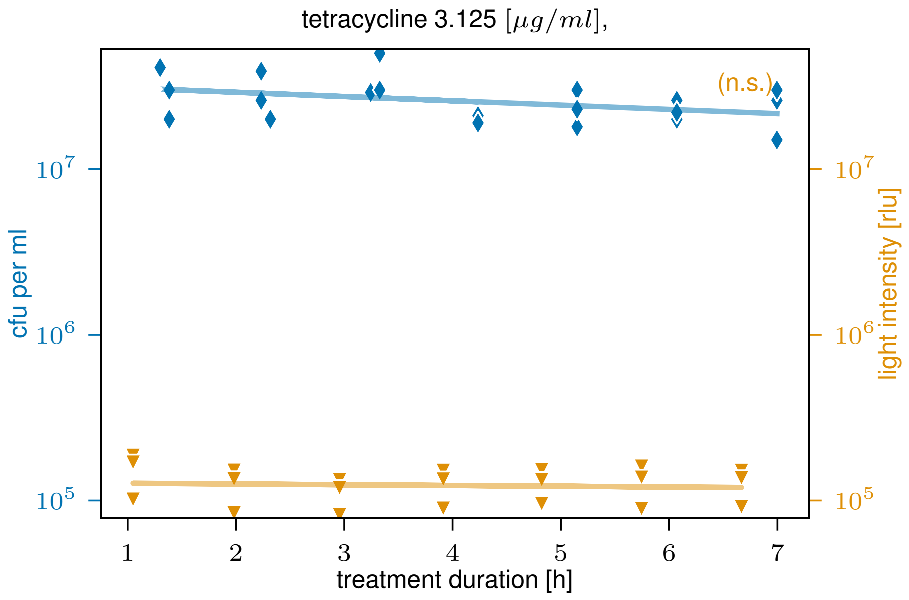


**Figure S18:** Tetracycline. The CFU signal is shown as blue diamonds and the light intensity as orange triangles for data points above their respective detection limits ($10^4$ CFU/mL for CFU and 20 rlu for luminescence). Lines represent log-linear fits to the corresponding signals.

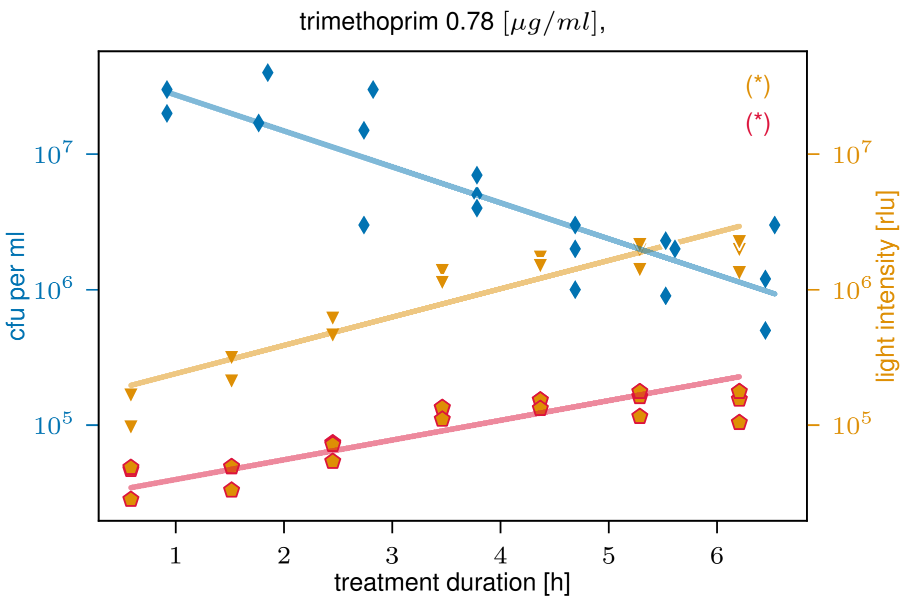


**Figure S19:** Trimethoprim. The CFU signal is shown as blue diamonds and the light intensity as orange triangles for data points above their respective detection limits ($10^4\,\mathrm{CFU/mL}$ for CFU and 20 rlu for luminescence). Lines represent log-linear fits to the corresponding signals. The morphology-corrected luminescence signal is shown as orange pentagons with red frames, and the corresponding rate fit indicated by a red line.

a)

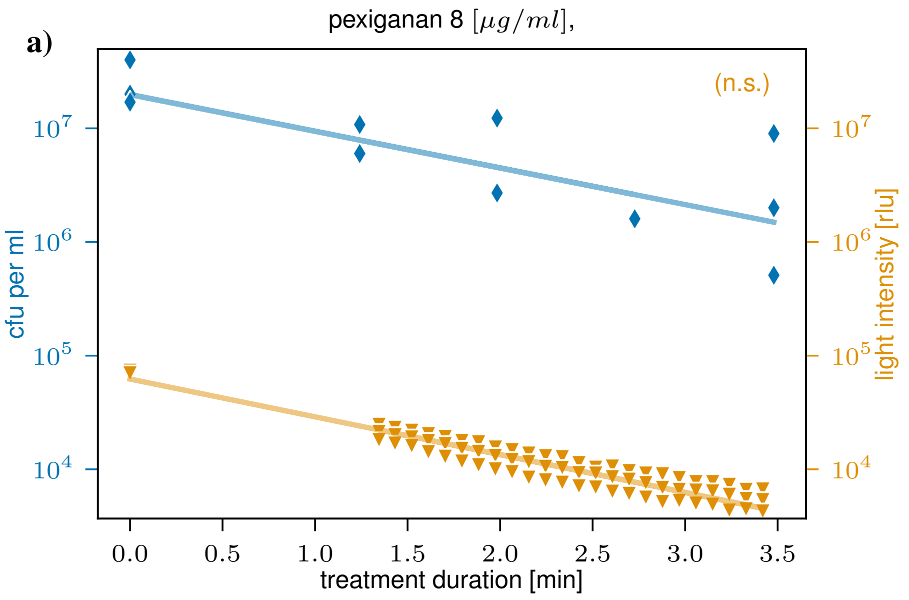


b)

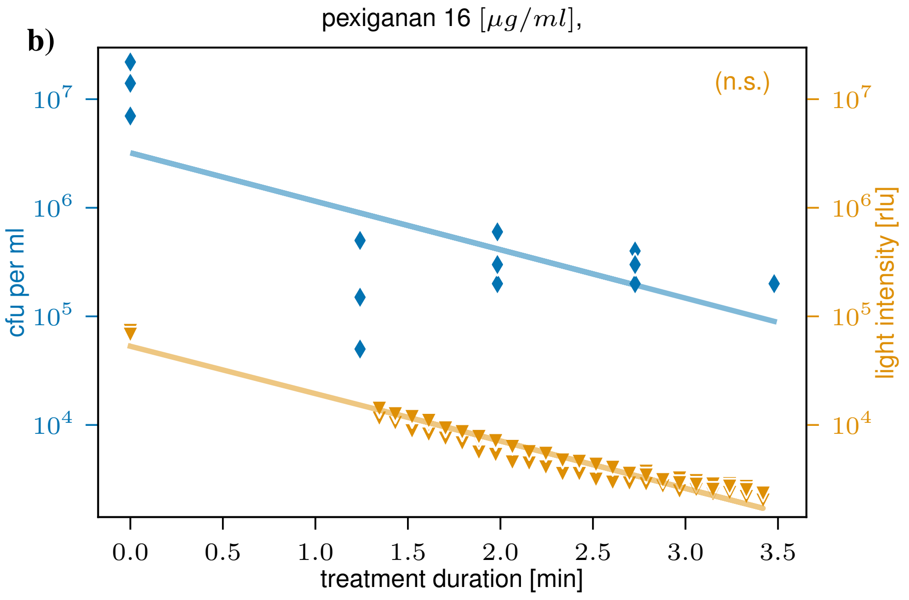


**Figure S20:** Comparison of the CFU and luminescence signals from the pexiganan kill curve experiment using the liquid handling platform. The first data points at $t_0$ represent the pretreatment CFU and light intensity values.

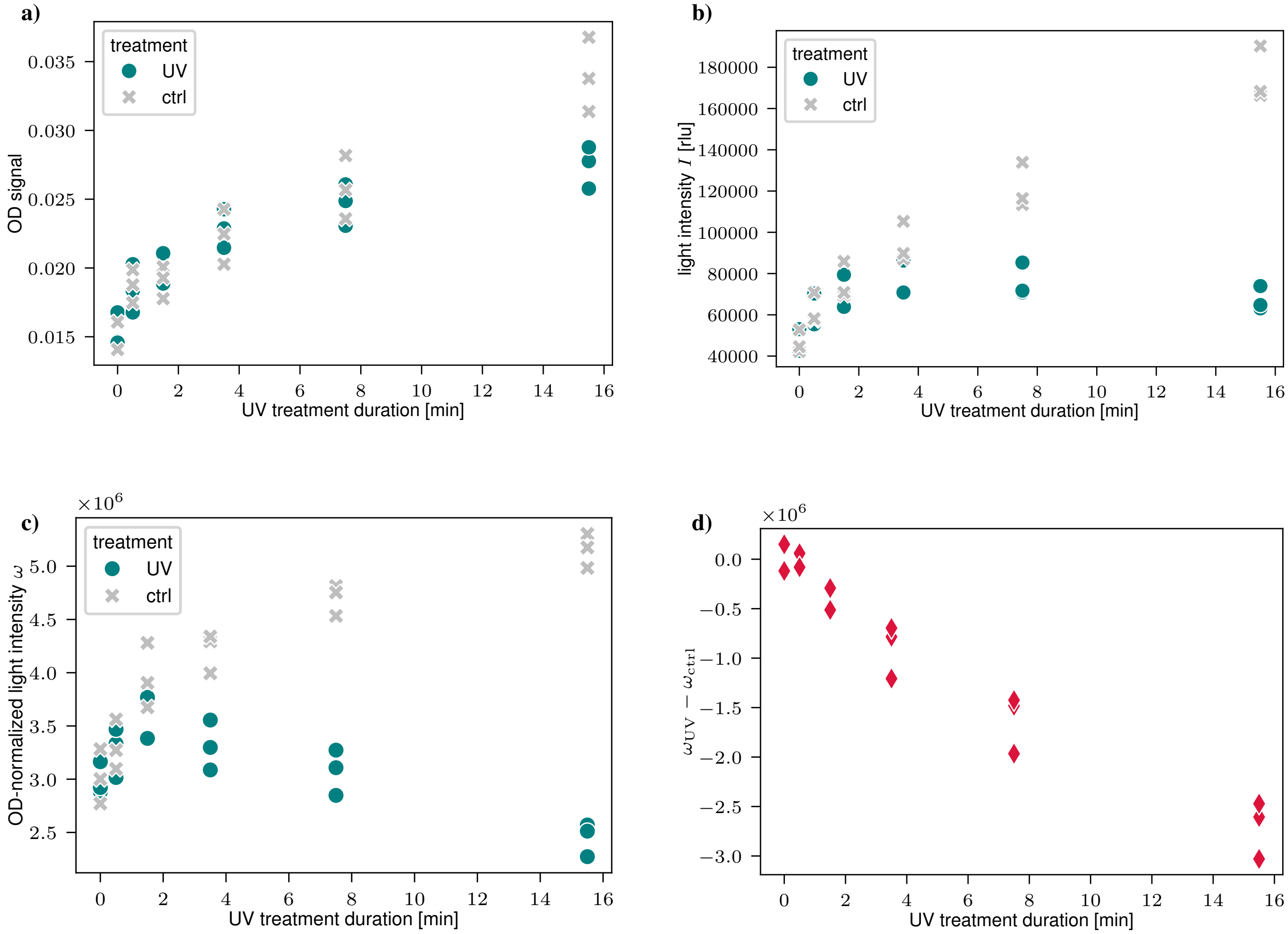


**Figure S21:** Panel plot showing the effects of UV treatment on bacterial density (approximated by OD) and light intensity (I) over time by comparing treated (UV) and untreated (ctrl) cultures. (a) Optical density (OD), (b) light intensity, (c) OD-specific light intensity $\omega = I(t)/\mathrm{OD}(t)$, and (d) the difference in OD-specific light intensity between UV-treated and control.

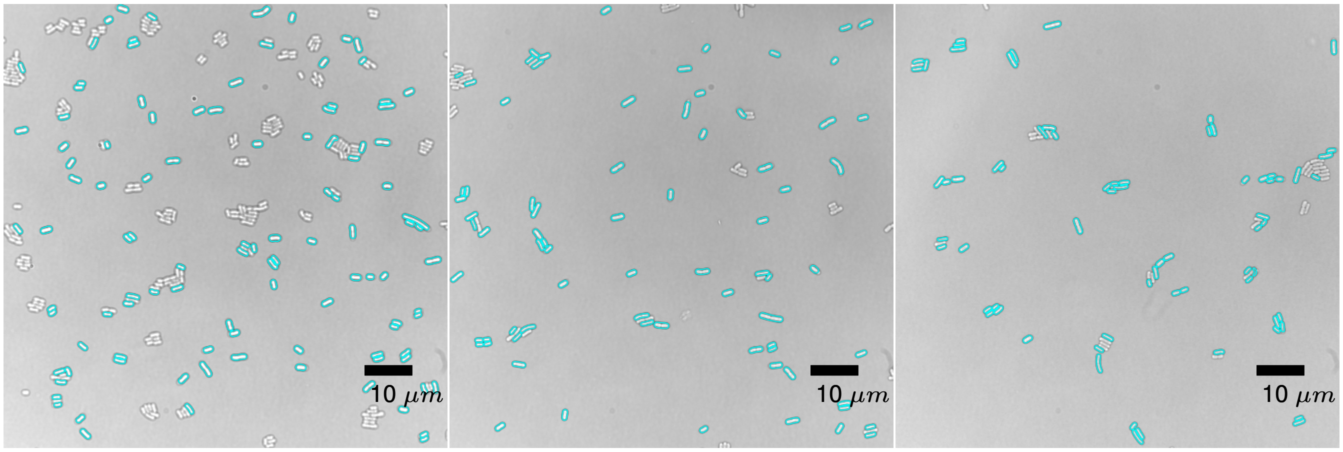


**Figure S22:** Microscopy images of control (before treatment) samples. Each image represents a different replicate. We plotted the green channel of the recorded images in greyscale. The red channel, capturing propidium iodide activity, is overlaid in red. Bacterial shapes detected by the algorithm are outlined in cyan.

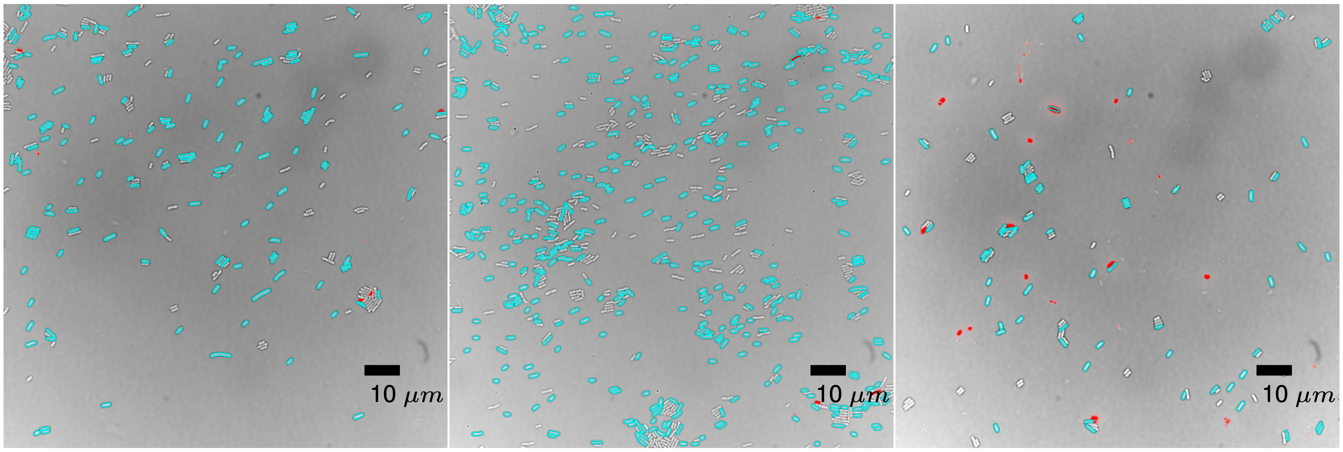


**Figure S23:** Microscopy images of control (after 2 hours) samples. Each image represents a different replicate. We plotted the green channel of the recorded images in greyscale. The red channel, capturing propidium iodide activity, is overlaid in red. Bacterial shapes detected by the algorithm are outlined in cyan.

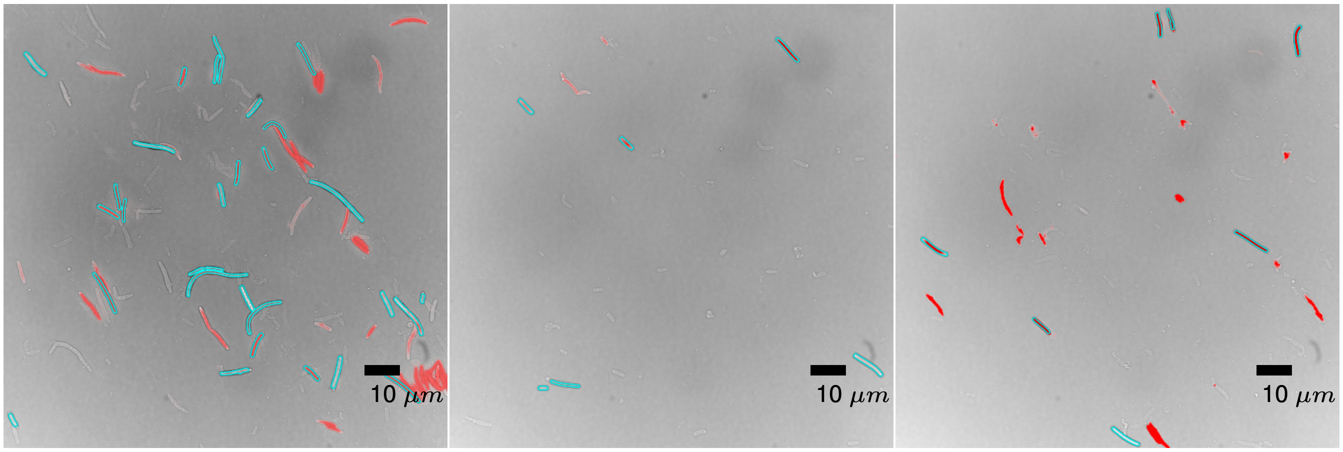


**Figure S24:** Microscopy images of cells treated with ampicillin. Each image represents a different replicate. We plotted the green channel of the recorded images in greyscale. The red channel, capturing propidium iodide activity, is overlaid in red. Bacterial shapes detected by the algorithm are outlined in cyan.

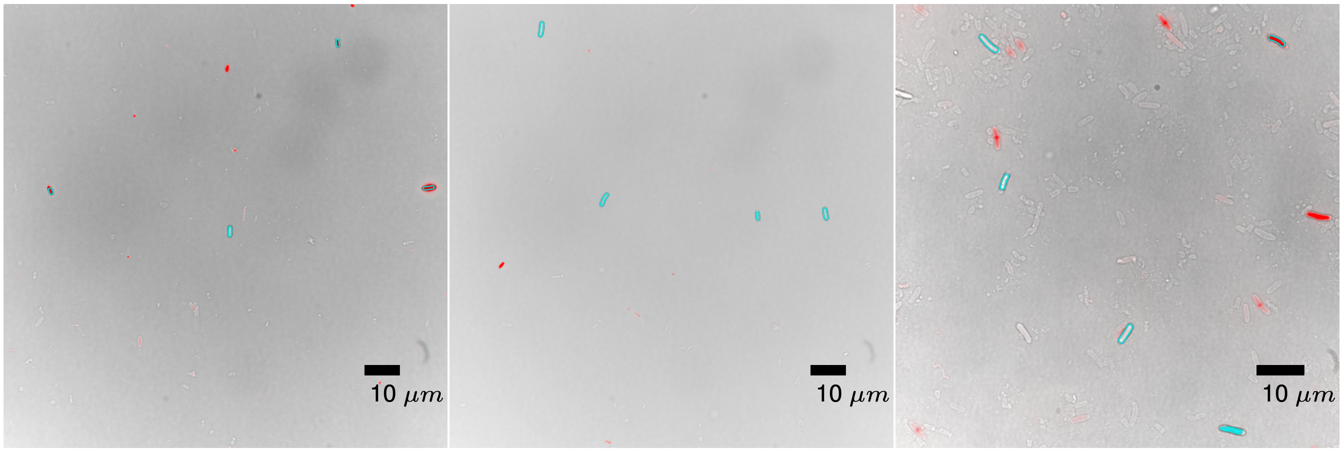


**Figure S25:** Microscopy images of cells treated with amoxicillin. Each image represents a different replicate. We plotted the green channel of the recorded images in greyscale. The red channel, capturing propidium iodide activity, is overlaid in red. Bacterial shapes detected by the algorithm are outlined in cyan.

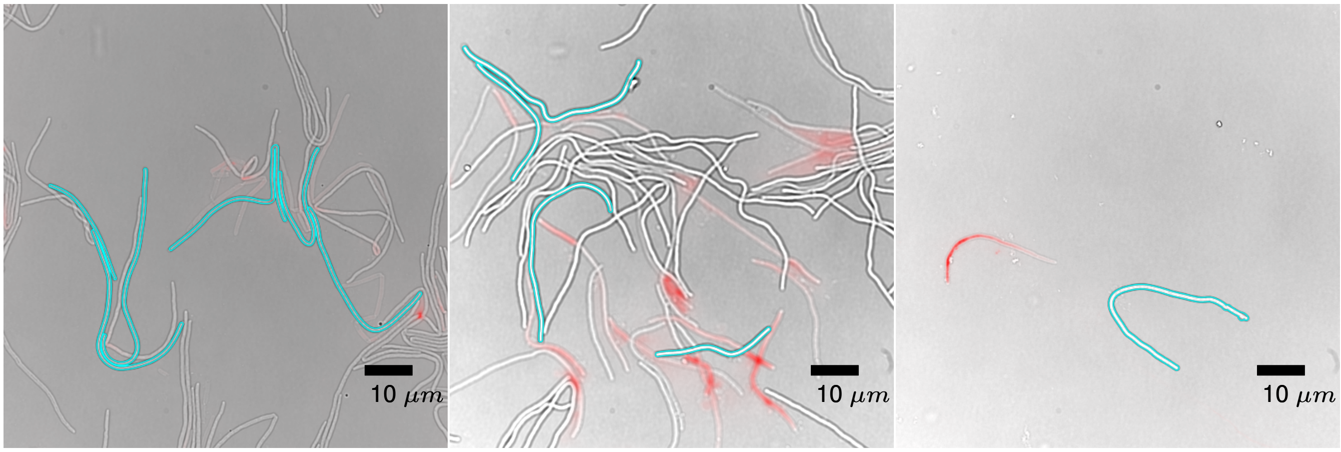


**Figure S26:** Microscopy images of cells treated with ceftazidime. Each image represents a different replicate. We plotted the green channel of the recorded images in greyscale. The red channel, capturing propidium iodide activity, is overlaid in red. Bacterial shapes detected by the algorithm are outlined in cyan.

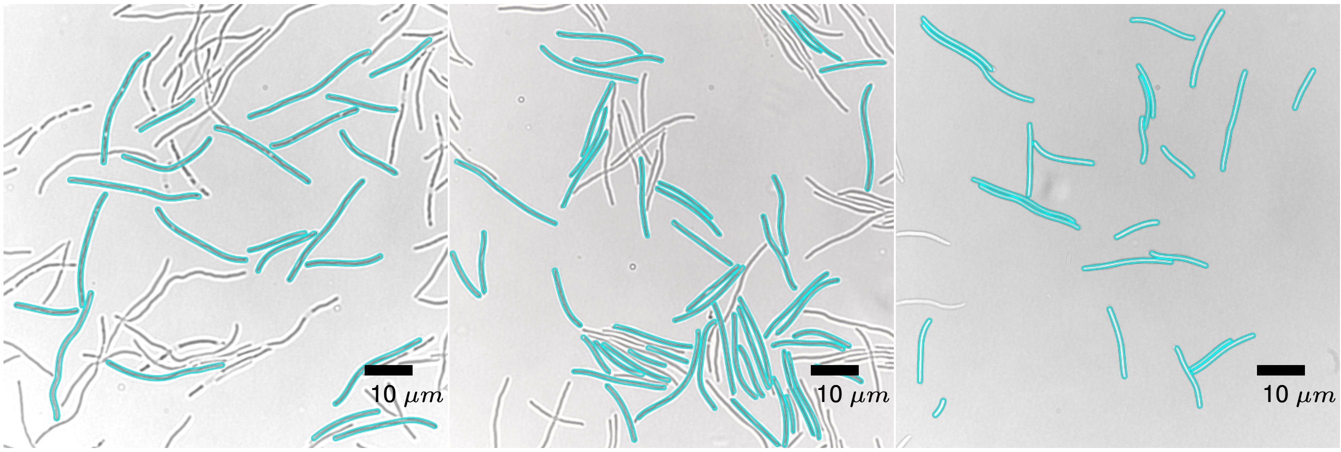


**Figure S27:** Microscopy images of cells treated with ciprofloxacin. Each image represents a different replicate. The white gaps can indicate the start of cell division (cells are not fixed). We plotted the green channel of the recorded images in greyscale. The red channel, capturing propidium iodide activity, is overlaid in red. Bacterial shapes detected by the algorithm are outlined in cyan.

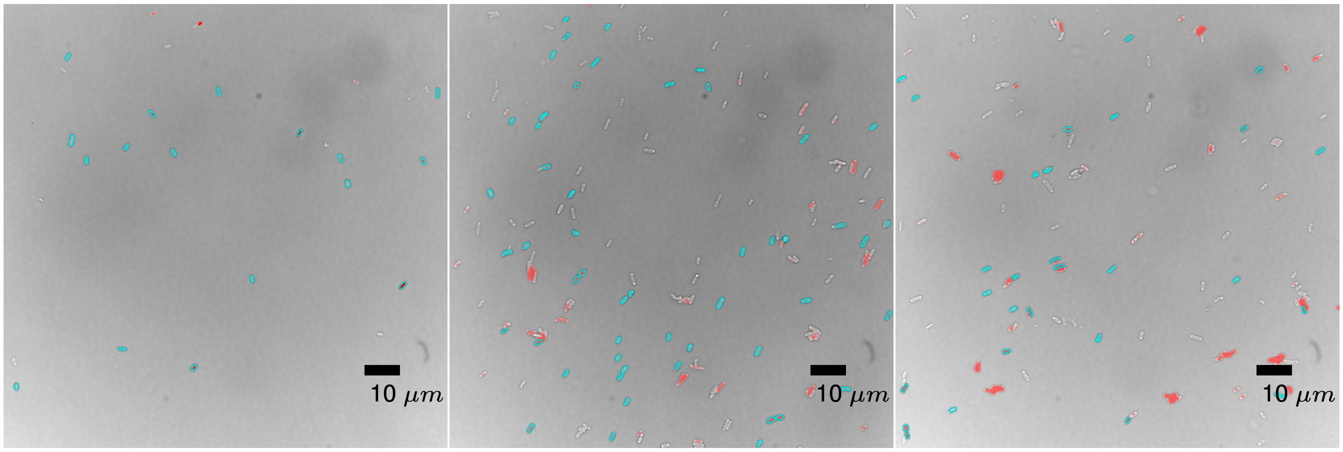


**Figure S28:** Microscopy images of cells treated with colistin. Each image represents a different replicate. We plotted the green channel of the recorded images in greyscale. The red channel, capturing propidium iodide activity, is overlaid in red. Bacterial shapes detected by the algorithm are outlined in cyan.

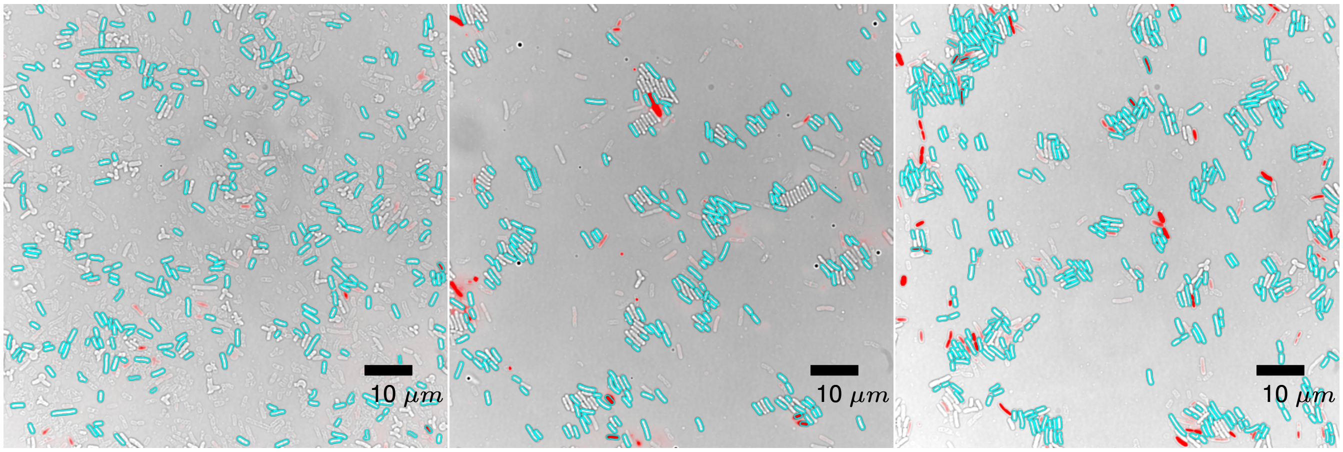


**Figure S29:** Microscopy images of cells treated with fosfomycin. Each image represents a different replicate. We plotted the green channel of the recorded images in greyscale. The red channel, capturing propidium iodide activity, is overlaid in red. Bacterial shapes detected by the algorithm are outlined in cyan.

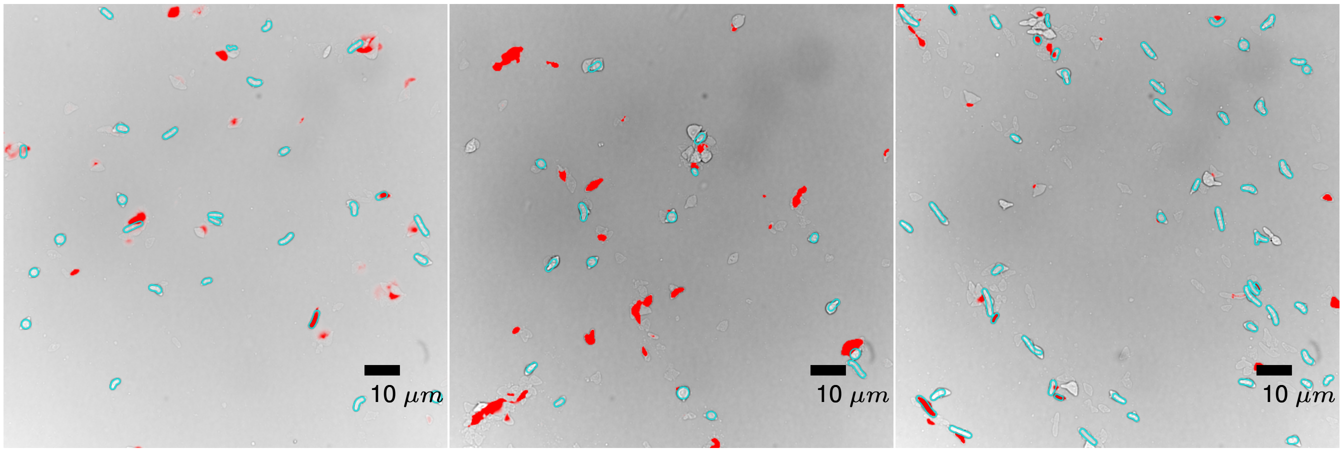


**Figure S30:** Microscopy images of cells treated with meropenem. Each image represents a different replicate. We plotted the green channel of the recorded images in greyscale. The red channel, capturing propidium iodide activity, is overlaid in red. Bacterial shapes detected by the algorithm are outlined in cyan.

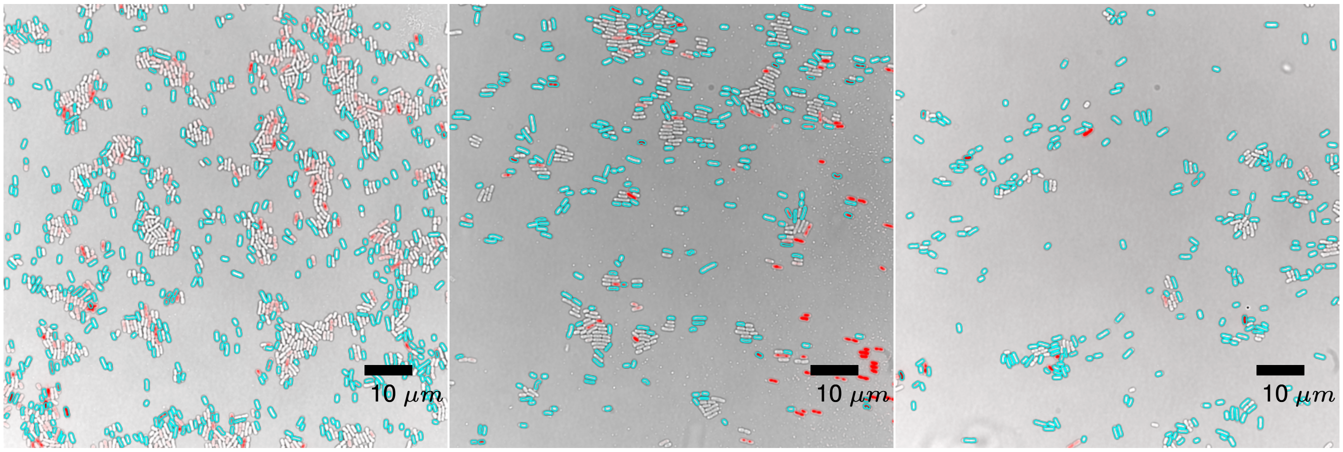


**Figure S31:** Microscopy images of cells treated with pexiganan. Each image represents a different replicate. We plotted the green channel of the recorded images in greyscale. The red channel, capturing propidium iodide activity, is overlaid in red. Bacterial shapes detected by the algorithm are outlined in cyan.

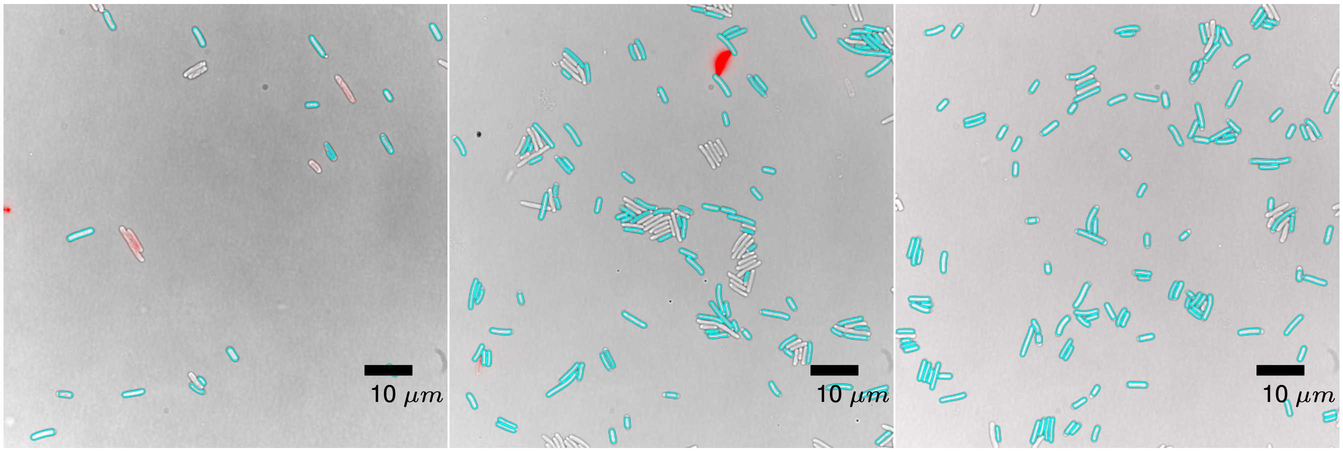


**Figure S32:** Microscopy images of cells treated with rifampicin. Each image represents a different replicate. We plotted the green channel of the recorded images in greyscale. The red channel, capturing propidium iodide activity, is overlaid in red. Bacterial shapes detected by the algorithm are outlined in cyan.

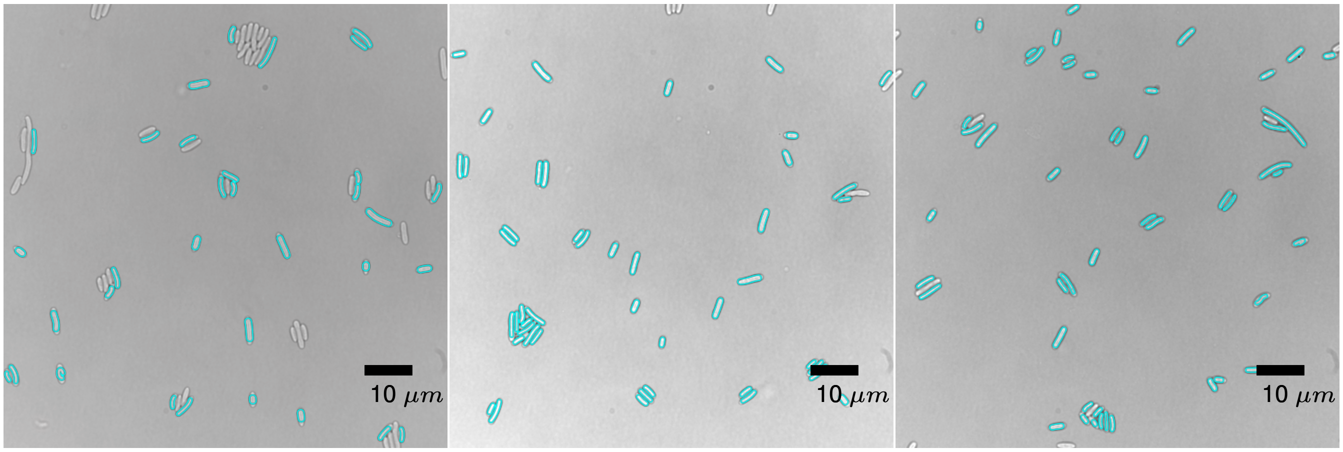


**Figure S33:** Microscopy images of cells treated with tetracycline. Each image represents a different replicate. We plotted the green channel of the recorded images in greyscale. The red channel, capturing propidium iodide activity, is overlaid in red. Bacterial shapes detected by the algorithm are outlined in cyan.

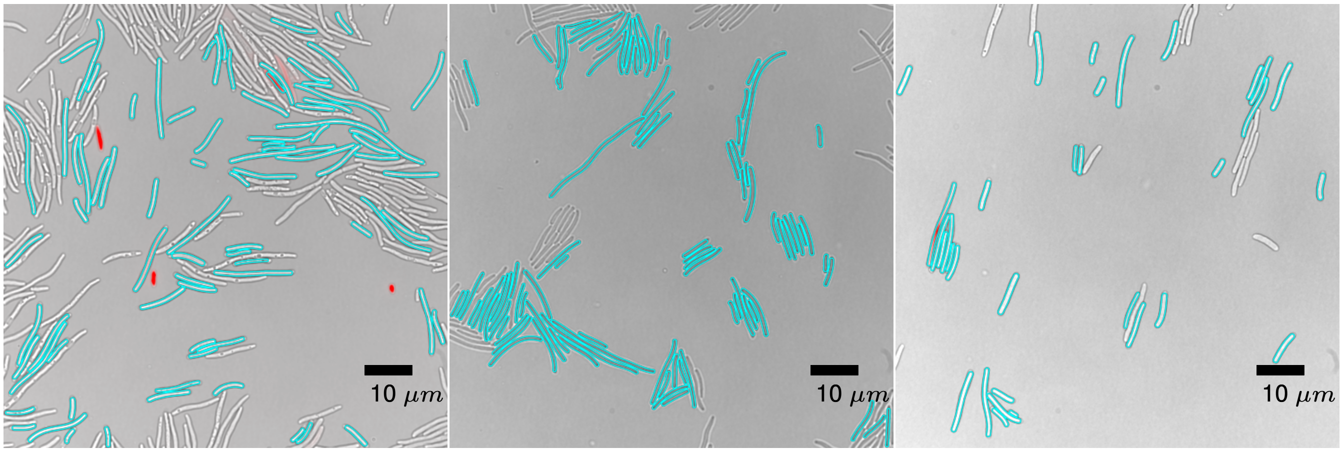


**Figure S34:** Microscopy images of cells treated with trimethoprim. Each image represents a different replicate. We plotted the green channel of the recorded images in greyscale. The red channel, capturing propidium iodide activity, is overlaid in red. Bacterial shapes detected by the algorithm are outlined in cyan.

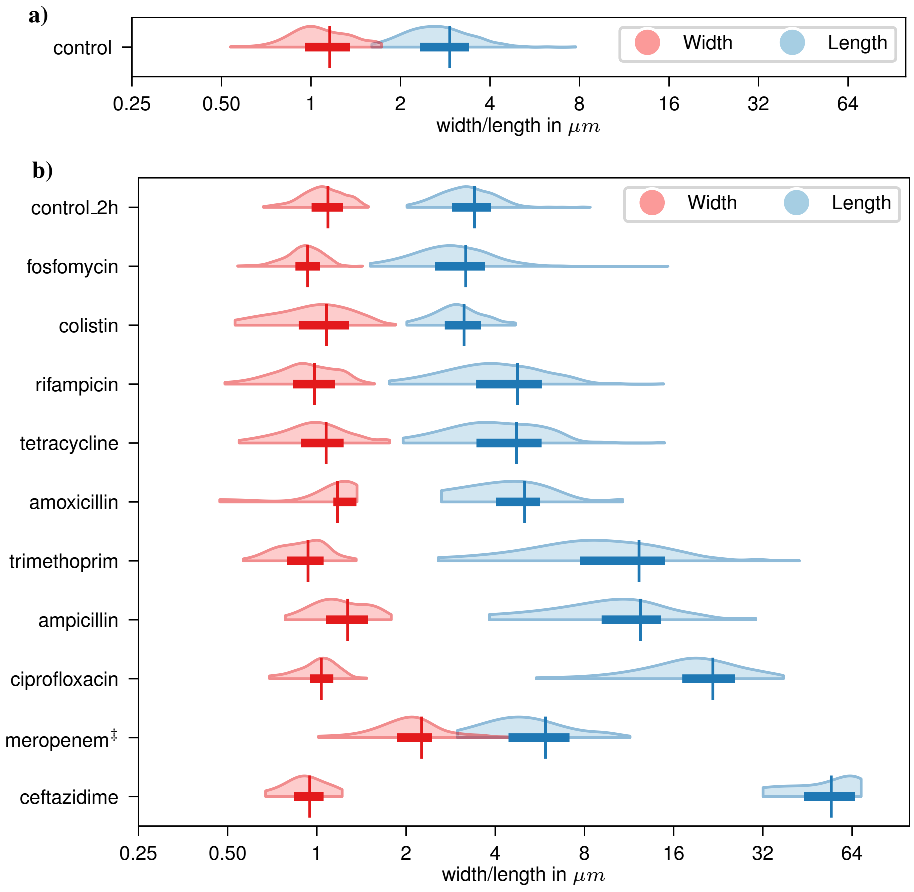


**Figure S35:** Pooled density distributions (95% prediction intervals) of cell widths (red) and lengths (blue), obtained from microscopy images (a) before and (b) after 2 h of antibiotic treatment. Boxes indicate the interquartile range (Q1–Q3), and the mean is marked by (|). ‡ Poor fit quality for meropenem-treated cells, which adopt a lemon-like shape (Figure S30), due to our algorithm assuming cylindrical geometry.

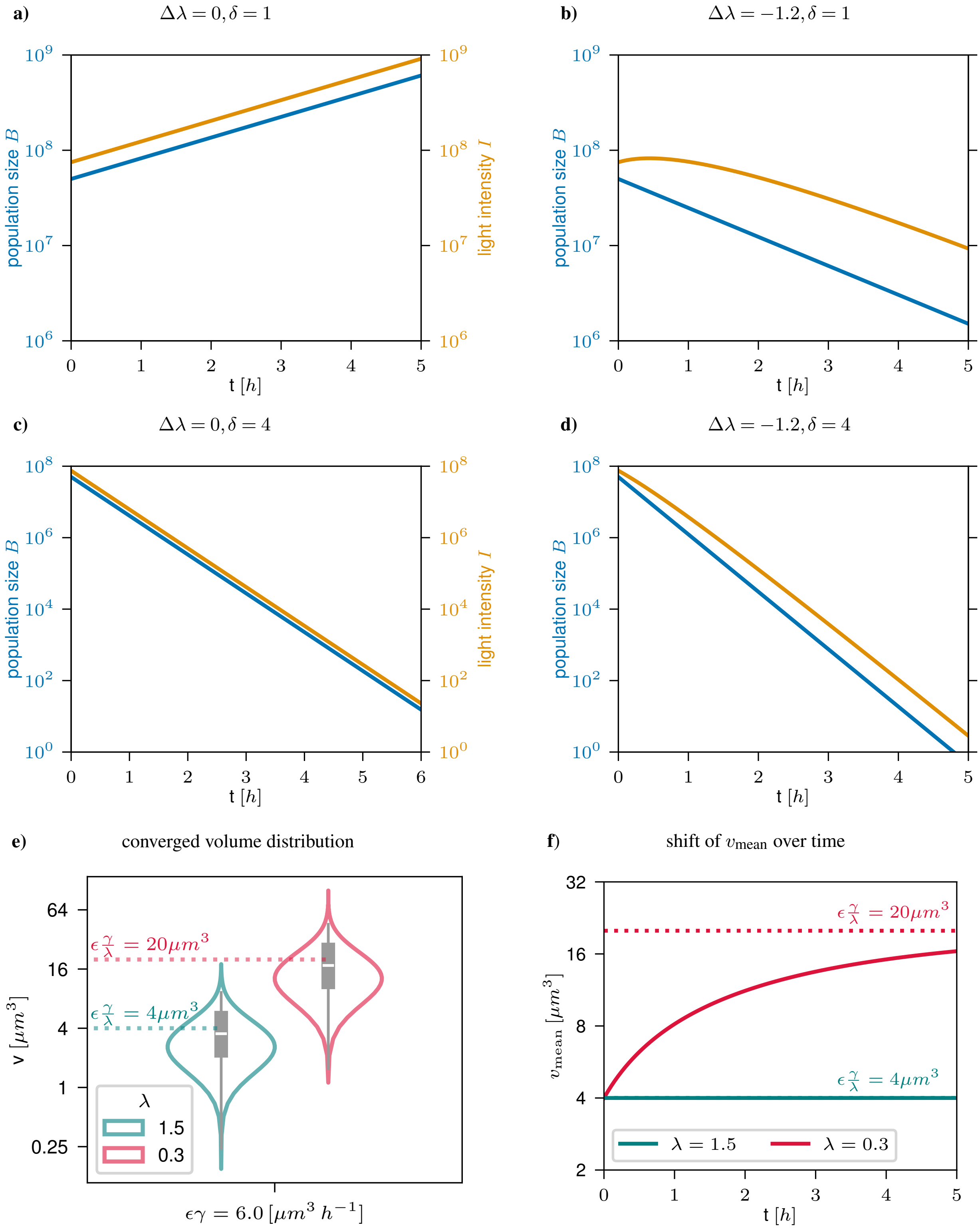


**Figure S36:** Illustrative simulations using the filamentation model relating (a–d) bacterial population size (blue) and light intensity (orange) under different combinations of treatment-induced changes in division rate ($\Delta\lambda$) and death rate ($\delta$). Panel (e) shows the distributions of converged cell volumes for $\lambda = 0.3$ and $\lambda = 1.5$. Panel (f) shows the shift of mean cell volumes over time for $\Delta\lambda = -1.2$ and $\Delta\lambda = 0$. For all simulations, we used $\lambda_0 = 1.5$, $\gamma = 150$, $\phi = 0.015$, and $\epsilon = 0.04$. As shown in panel (b), treatment-induced filamentation can lead to a temporary discrepancy between luminescence- and CFU-based rates.

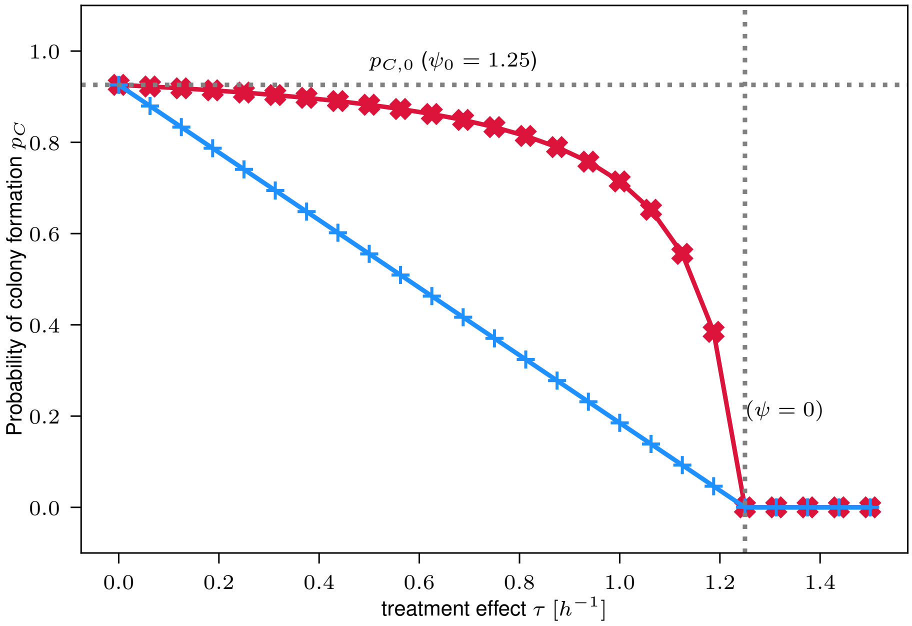


**Figure S37:** Example of the probability of colony formation $p_C = 1 - p_E$ (see Appendix S2) for a single plated bacterium. Blue shows $p_C$ for a purely bactericidal drug ($\lambda_T = 0$) and red for a purely bacteriostatic drug ($\delta_T = 0$), plotted over the treatment effect $\tau = \delta_T + \lambda_T$. In this illustrative example we use $\lambda_0 = 1.35h^{-1}$ and $\delta_0 = 0.1h^{-1}$.

**a)**

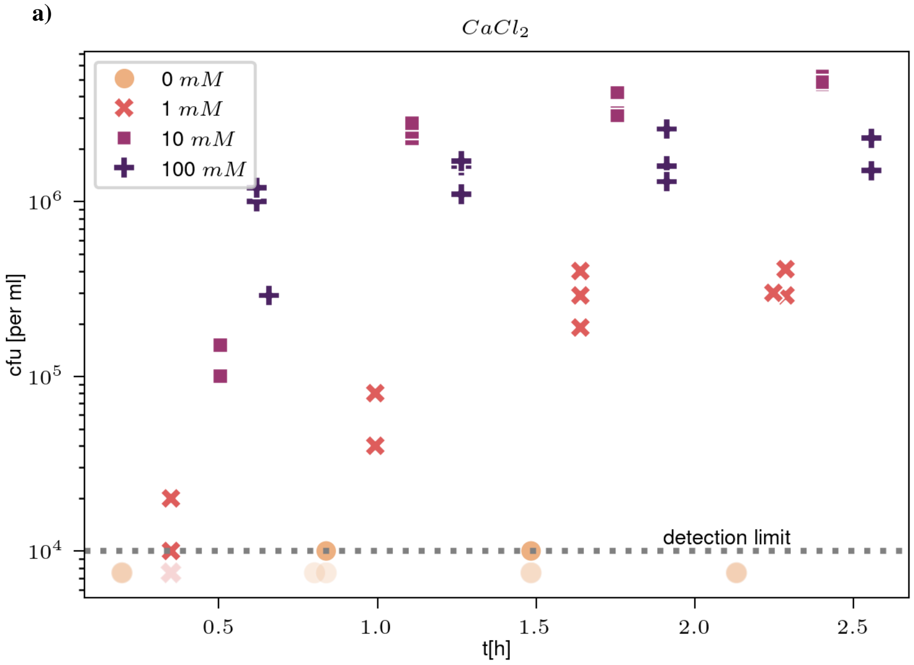


**b)**

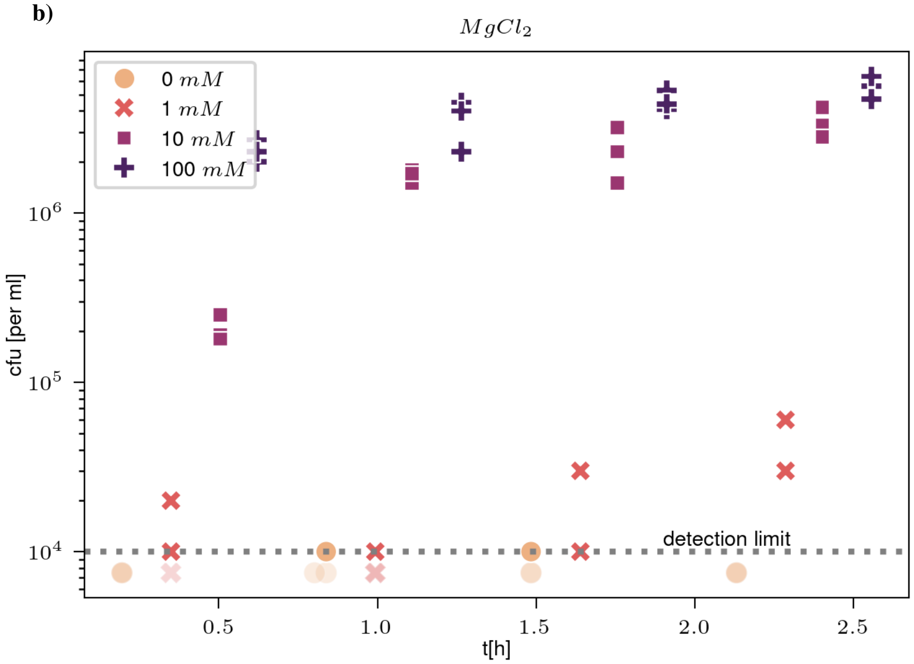


**Figure S38:** CFU measured over time in supplemented dilution media. Bacterial cultures treated for 1 min with $16\,\mu\mathrm{g/ml}$ pexiganan were diluted (1:100) in PBS supplemented with varying concentrations of (a) $CaCl_2$ and (b) $MgCl_2$. Diluted samples were repeatedly plated over time to test whether supplementation prevents further bacterial killing.

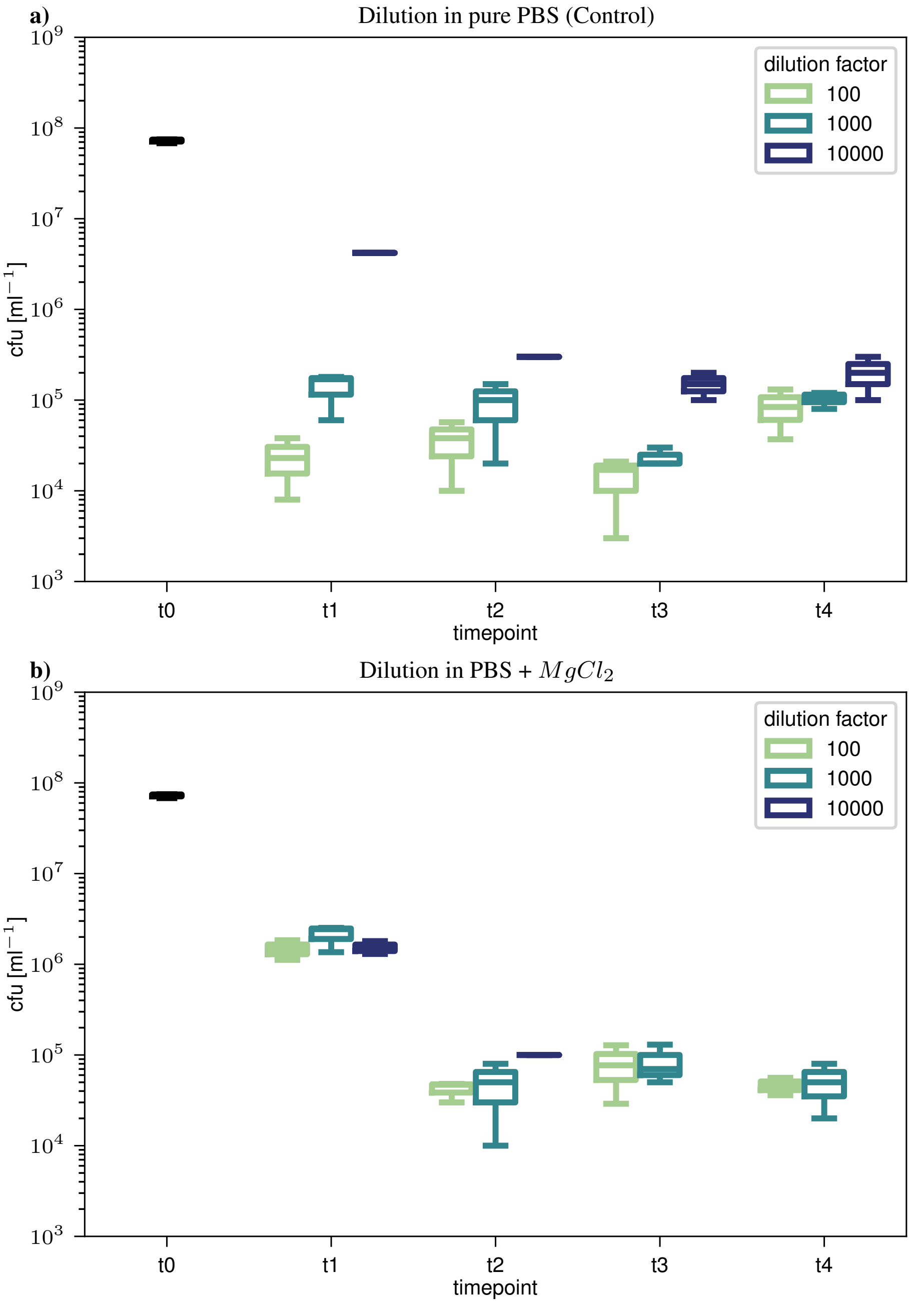


**Figure S39:** CFU time-kill curves for pexiganan, performed manually, comparing dilution in (a) unsupplemented PBS and (b) PBS supplemented with $100\,\mathrm{mM}$ $MgCl_2$. Timepoints correspond to: $t_0 \approx 0\,\mathrm{s}$, $t_1 \approx 20\,\mathrm{s}$, $t_2 \approx 2\,\mathrm{min}$, $t_3 \approx 3\,\mathrm{min}\,20\,\mathrm{s}$, and $t_4 \approx 5\,\mathrm{min}$. The black boxplot ($t_0$) indicates the pre-treatment bacterial density.

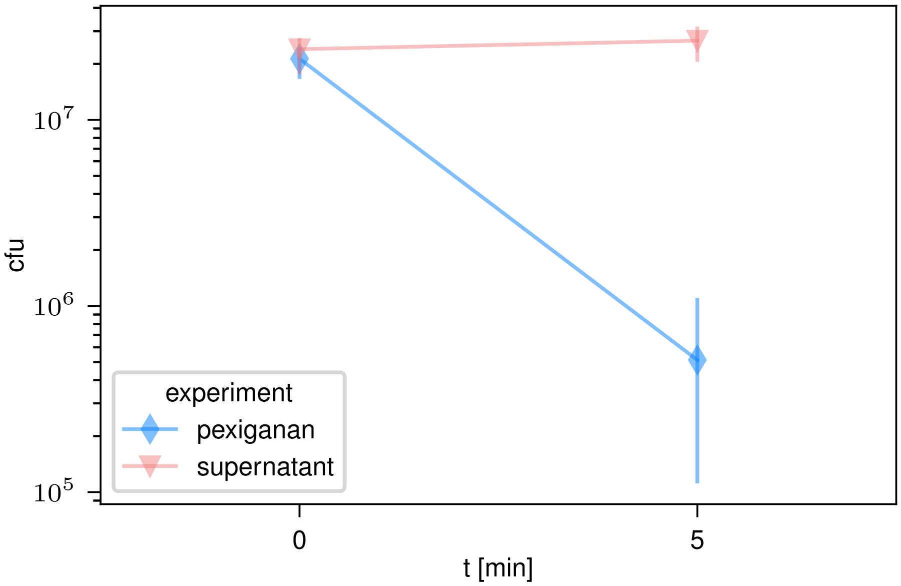


**Figure S40:** Experiment AMP kill curve vs supernatant kill curve. Here we show the original AMP-Killcurve in blue and the change of CFU over time in the supernatant. The error bars show the min/max interval of the three replicates. Table S5 lists the confidence interval and mean for the bootstrapped rates. According to the significance criterion (defined in methods) these two rates are significantly different. The confidence interval of the rate of change of CFU in the supernatant includes zero.
